# The type of inhibition provided by thalamic interneurons alters the input selectivity of thalamocortical neurons

**DOI:** 10.1101/2024.01.17.576001

**Authors:** Deyl Djama, Florian Zirpel, Zhiwen Ye, Gerald Moore, Charmaine Chue, Christopher Edge, Polona Jager, Alessio Delogu, Stephen G Brickley

## Abstract

A fundamental problem in neuroscience is how neurons select for their many inputs. A common assumption is that a neuron’s selectivity is largely explained by differences in excitatory synaptic input weightings. Here we describe another solution to this important problem. We show that within the first order visual thalamus, the type of inhibition provided by thalamic interneurons has the potential to alter the input selectivity of thalamocortical neurons. To do this, we developed conductance injection protocols to compare how different types of synchronous and asynchronous GABA release influence thalamocortical excitability in response to realistic patterns of retinal ganglion cell input. We show that the asynchronous GABA release associated with tonic inhibition is particularly efficient at maintaining information content, ensuring that thalamocortical neurons can distinguish between their inputs. We propose a model where alterations in GABA release properties results in rapid changes in input selectivity without requiring structural changes in the network.

## Introduction

Understanding how an individual neuron extracts information from the complex patterns of synaptic input arriving onto their dendrites is fundamental to our understanding of brain function. In this study, we focused on thalamocortical (TC) neurons of the dorsal lateral geniculate nucleus (LGd) where many of the rules governing input selection have been developed. For example, early simultaneous recordings between retinal ganglion cells (RGCs) and TC neurons in the cat LGd first described how a single neuron was excited by one or two RGC-types (Bishop *et al*., 1958; Cleland *et al*., 1971) and this *in vivo* observation has held true in other mammals including the mouse (Grubb & Thompson, 2003). However, high-resolution anatomical reconstruction techniques have described the convergence of multiple retinogeniculate excitatory synapses onto a single TC neuron (Hammer *et al*., 2015; Morgan *et al*., 2016) and rabies virus tracing techniques have confirmed that multiple RGC-types often converge onto a single TC neuron (Rompani *et al*., 2017). This is often referred to as the “fuzzy logic” problem. Appreciation of this anatomical complexity has further highlighted the importance of synaptic plasticity mechanisms for generating input preference (Usrey *et al*., 1999; Litvina & Chen, 2017) leading to the idea that axonal inputs associated with a greater synaptic strength transmit the most salient information. In this study, we explored whether the inhibition generated by LGd interneurons (Jager *et al*., 2016; Morgan & Lichtman, 2020; Bakken *et al*., 2021; Jager *et al*., 2021) provides an additional level of control capable of altering TC neuron responses to RGC input.

GABA signaling within the mammalian thalamus reflects release from several sources. Axonal GABA release from the thalamic reticular nucleus (nRT) (Sumitomo *et al*., 1976; Montero & Scott, 1981) provides conventional axo-somatic and axo-dendritic inhibition at so-called F1-type synapses. Local interneuron mediated GABA release occurs both at conventional F1-type axonal synapses and the more unusual dendro-dendritic or F2-type synapses (Ohara *et al*., 1983; Ferraguti *et al*., 2004) that are thought to be restricted to the triadic glomerular synapses identified on TC neuron dendrites (Sherman, 2004). Finally, the steady-state dynamics of the GABA transporter largely determines the resting GABA concentration (Wu *et al*., 2007) that is responsible for the persistent activation of high-affinity extra synaptic GABA_A_ receptors and the resulting generation of a tonic conductance in TC neurons (Bright *et al*., 2011; Brickley & Mody, 2012; Jager *et al*., 2016). Although these distinct types of vesicular and non-vesicular GABA release mechanisms have been described in many different brain regions (Brickley & Mody, 2012), little is known about the role these distinct and unique types of inhibitory conductance changes play in the transfer of sensory information within the thalamus.

The dynamic-clamp protocols we developed to study the impact of these F1-type, F2-type and tonic forms of inhibition, enabled a single TC neuron to be excited with 10 different RGC input patterns while allowing the synaptic weight at each of these inputs to be varied. The synaptic conductance waveforms simulated the convergence of several excitatory inputs onto a single TC neuron based upon RGC recordings obtained from the mouse retina during stimulation with realistic visual scenes (Meytlis *et al*., 2012). A similar methodology was used to define the most energy efficient synaptic weighting at the retinogeniculate synapse (Harris *et al*., 2015). We expanded this approach to simultaneously mimic feedforward inhibition onto TC neurons based upon the scenario that local LGd interneurons were receiving excitation from the same RGC inputs that excited the TC neurons (Morgan & Lichtman, 2020). The properties of F1-type and tonic inhibitory conductance changes have been well described by many laboratories including our own (Bright *et al*., 2007). Optogenetics targeted to Sox14 expressing LGd interneurons was combined with other anatomical and functional approaches to explore the characteristics of dendro-dendritic GABA release at the glomerular F2-type synapse formed by LGd interneurons. This information was used to design novel dynamic-clamp protocols that explored the impact of F1-type, F2-type and tonic inhibition on a TC neurons response to retinogeniculate input. The level of control afforded by this approach enables a quantitative assessment of features such as TC neuron input sensitivity, AP output precision, rate coding, gain control and information loss in the face of changing levels of feedforward inhibition. The observations made in this study lead us to propose a novel model for controlling input selectivity that offers computational flexibility with low metabolic demand that should be applicable to other brain regions that rely on both vesicular and non-vesicular sources of GABA release.

## Materials and Methods

### Mouse strains and viral delivery

Most experiments used C57Bl6 mice. Sox14gfp/+ and Sox14cre/+ mice have the Sox14 coding sequence replaced by the eGFP gene (Crone *et al*., 2008; Delogu *et al*., 2012) and Cre gene (Jager *et al*., 2021; Brock *et al*., 2022) respectively. A single copy of the wild type Sox14 allele is retained in Sox14gfp/+ and Sox14cre/+ mice and this is sufficient to maintain gene function. Heterozygote Sox14gfp/+ animals display a normal number of thalamic interneurons (see (Jager *et al*., 2021)) with no overt behavioural or morphological phenotype. The nomenclature Sox14gfp/+ and Sox14cre/+ indicates that one copy of the wild type Sox14 allele is retained in these mice. These mice are not Sox14 loss-of-function animals. The nomenclature Sox14gfp/gfp and Sox14cre/cre indicates that both copies of Sox14 allele are modified and, therefore, these are Sox14 loss-of-function mutant animals (Jager *et al*., 2021).

Adeno-associated virus (AAV) type 2 carrying cre-dependent ChR2-eYFP (Addgene plasmid 20298, pAAV-EF1a−double floxed−hChR2 (H134R)-EYFP-WPRE-HGHpA) were injected at postnatal day 16 (P16). 0.5µl of this AAV virus was bilaterally injected into the lateral geniculate nucleus 2 weeks before electrophysiological experiments were performed.

### Electrophysiology

Acute brain slices were prepared from adult mice that were killed by cervical dislocation (in accordance with UK Home Office guidelines). The brain was rapidly removed and immersed in ice cold slicing solution. The slicing solution contained (in mM: NMDG 92, KCl 2.5, NaH2PO4 1.25, 30 mM NaHCO3, HEPES 20, glucose 25, thiourea 2, Na-ascorbate 5, Na-pyruvate 3, CaCl2·4H2O 0.5 and MgSO4·7H2O 10). The pH was adjusted to 7.3 with concentrated HCl when bubbled with 95%O_2_/5%CO_2_. Slices were prepared on a vibrating microtome (7000smz-2, Campden instruments Ltd., Loughborough, UK) at a thickness of 300 µm and immediately transferred to a holding chamber containing slicing ACSF continuously bubbled with 95%O_2_/5%CO_2_. Once slicing was complete, the holding chamber was transferred to a 37°C heat block for 40 minutes after which the slicing ACSF was gradually exchanged for recording ACSF (in mM: NaCl 125, KCl 2.5, CaCl_2_ 2, MgCl 2, NaH_2_PO4 1.25, NaHCO_3_ 26, glucose 11, pH 7.4 when bubbled with 95%O_2_/5%CO_2_) and allowed to reach room temperature whilst the solutions were exchanged prior to electrophysiological recording.

Slices were visualized with infra-red illumination (M780LP1 LED, Thorlabs Ltd., Ely, UK) on fixed-stage upright microscopes (BX51W1, Olympus Ltd., Tokyo, Japan or Axioscop 2, Zeiss Ltd., Jena, Germany) fitted with high numerical aperture water-immersion objectives and digital cameras (KP-M1AP, Hitachi Europe Ltd., Maidenhead, UK). Patch pipettes were fabricated from thick-walled borosilicate glass capillaries (GC150F, Harvard Apparatus Inc., Cambridge, UK) using a two-step vertical puller (PC-10, Narishige International Ltd., London, UK). Pipette resistances were typically 3-4 MΩ when back filled with internal solution. For voltage-clamp experiments, the internal solution contained (in mM: CsCl 140, NaCl 4, CaCl_2_ 0.5, HEPES 10, EGTA 5, Mg-ATP 2; the pH was adjusted to 7.3 with CsOH). Biocytin (1.5 mg/ml) was included in the pipette solution so that neuronal cell-type could be confirmed. The command voltage for all voltage-clamp experiments was −70mV and we did not correct for the liquid junction potential of −9.8 mV that was present with the internal and external solutions. Unless otherwise stated, the broad-spectrum glutamate receptor antagonist kynurenic acid at a 100 μM concentration was added to the external solution during voltage-clamp experiments to record sIPSCS in isolation. In some experiments the broad spectrum GABA_A_ receptor antagonist gabazine (1 μM) was added to the external solution at the end of the experiment to record the properties of sEPSCs in isolation and quantify the level of tonic inhibition. For current-clamp recording the internal solution contained (in mM: C_6_H_11_KO_7_ 140, NaCl 4, KCl 5, HEPES 10, EGTA 5, Mg-ATP 4, Na-ATP 0.3, CaCl_2_ 0.5; the pH was adjusted to 7.3 with KOH). Biocytin (1.5 mg/ml) was also included in the pipette solution. Pipette placement was controlled by micromanipulators (PatchStar, Scientifica Ltd, Uckfield, UK) that mounted upon a motorized stage (UMS, Scientifica Ltd, Uckfield, UK). For voltage-clamp experiments, the amplifier current (Axopatch 700B, Molecular Devices; Foster City, CA) was filtered at 10 kHz (–3 dB, 8-pole low-pass Bessel) and digitized at 20 kHz using a National Instruments digitization board (PCI-6052E; National Instruments, Texas, USA). Data acquisition was performed using Signal (Version 6.03; Cambridge Electronic Devices; Cambridge, UK), WINWCP (Version 4.1.2) and WINEDR (Version 3.0.9) kindly provided by John Dempster (John Dempster; University of Strathclyde, UK). For current-clamp experiments, the voltage output of the amplifier was connected to a Power 1401 Series 3A (Cambridge Electronic Devices; Cambridge, UK) and was filtered at 10 kHz (–3 dB, 8-pole low-pass Bessel) before being sampled at 50 kHz. No additional steady state current was injected into the cell during current-clamp recording to influence the RMP.

For optogenetic experiments, a blue (470 nm) collimated LED (M470L3-C1, Thorlabs) was mounted to the back of the microscope and focused through the objective lens. The optical power emitted by our ×63 water-immersion lens increased linearly to a maximum power of 70□LμW□Lmm^−2^ at 1□LV. A 1 ms, 1□LV brief pulse protocol gave rise to a transient response that peaked at 40□LμWL□mm^−2^ with a 10–90% rise time of 0.73□Lms and a decay constant of 9.65□Lms; as determined from a single exponential fit. Optogenetic experiments were perforemed

### Serial Two-Photon Imaging

Sox14gfp/+ and Sox14gfp/gfp mouse brain samples were embedded in a 4.5% oxidised-agarose solution containing agarose (type 1; Sigma), 10 mM NaIO4 (Sigma) and 50 mM phosphate buffer (PB). Samples were imaged with TissueCyte 1000 (Ragan *et al*., 2012) with a custom cooling system (JULABO UK Ltd.) for serial two-photon (STP) tomography. Physical sectioning was performed every 50 μm with optical sectioning every 10 μm. A 16×, 0.8 NA immersion objective (Nikon Inc) acquired 1 × 1 mm image tiles at spatial resolution 0.54 μm with a 12 × 10 tiling mosaic required to obtain a complete coronal tissue section. Laser (Chameleon Ultra II, Coherent) excitation was conducted at 920 nm for GFP excitation with three PMT channel acquisition for red, green, and blue wavelength collection. Tiled data was stitched alongside STP acquisition using a custom Python and ImageJ/Fiji pipeline. STP data sets of each mouse brain were down-sampled to 10 μm isotropic voxel size and registered with the Allen CCF3 average atlas using Elastix (Klein et al., 2010). For automated cell counting, a U-Net deep learning network (Ronneberger O., 2015) was trained using Python and the TensorFlow 2.0 platform (Jager *et al*., 2021).

### Nearest Neighbour Distance Analysis

Nearest neighbour distance (NND) was obtained from 3D reconstructions of Sox14gfp/+ and Sox14gfp/gfp cell locations. The cells’ coordinates in 3D were generated by Neurolucida and analysed using a custom Python script and the Pandas library (McKinney et al., 2010) to calculate NNDs separately for each group and between the two groups, for each Sox14 brain individually. The data was then normalised to the largest NND within each data set (each individual group and between groups sets for each brain) averaged across the brains (mean ± SEM) and plotted as cumulative distribution. Normalisation allows us to plot their cumulative distribution as a fraction of the maximum distance, though even before normalisation of the curves were broadly similar. Statistically significant differences between the distributions were verified using the two-sample Kolmogorov–Smirnov test.

### Dynamic-clamp protocols

Dynamic-clamp protocols were run in Signal version 7.01 (Cambridge Electronic Devices; Cambridge, UK). The user-defined waveforms chosen to simulate time varying conductance changes at the retinogeniculate synapse were written in MathWorks (MATLAB, Cambridge, UK). The timing of each conductance change was taken from extracellular recordings of retinal ganglion cell (RGC) firing patterns recorded in response to a natural scene (Meytlis et al., 2012). The time varying conductance (*g(t)*) of each simulated retinogeniculate excitatory postsynaptic conductance (EPSC) was constructed using an exponential difference model of the type:

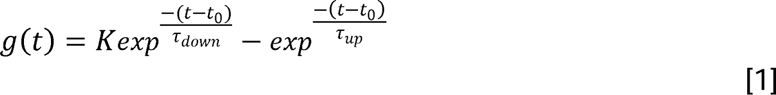

where K was a defined fraction of g_max_ (σ_max_). The properties of the exponential functions were chosen to mimc the presence of either AMPA-receptor mediated synaptic transmission EPSC_AMPA_ (τ_up_ = 0.1ms, τ_down_ = 3ms, σ_max_ = 1.0) or NMDA-receptor mediated synaptic transmission EPSC_NMDA_ (τ_up_ = 10ms, τ_down_ = 20ms, σ_max_ = 0.1). Additionally, synaptic dynamics at the retinogeniculate synapse was mimicked by convoluting the peak conductance of each EPSC_AMPA_ and EPSC_NMDA_ with a function inversely proportional to the instantaneous frequency of the RGC input pattern. The underlying I-V relationship of the EPSC_AMPA_ used to calculate the injected current (*I*_*inj*_) during dynamic-clamp was controlled by a linear leak of the type:

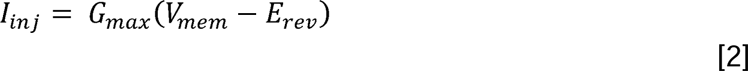

where *G*_*max*_ is the maximum conductance and V_mem_ is the membrane potential and E_rev_ is the reversal potential. In contrast, the IV relationship of the EPSC_NMDA_ was based upon a non-linear function of the type:

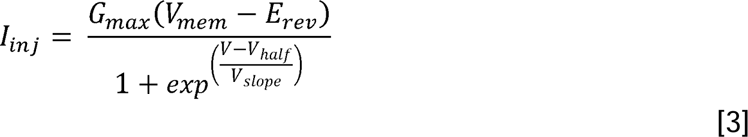

where *G*_*max*_ is the maximum conductance, *V_mem_* the membrane potential, *E_rev_* is the reversal potential, *V*_*half*_ is the potential at which 50% of NMDA channels are open in and *V_slope_* controls the range of the negative slope conductance. This relationship was chosen to mimic the presence of a voltage-dependent Mg^2+^ block of the NMDA receptor channels at the retinogeniculate synapse.

Three types of inhibition were simulated in the dynamic-clamp environment. Firstly, a tonic GABA_A_ receptor-mediated conductance (GABA_tonic_) was modelled with a time varying stochastic noise. Two distinct types of phasic inhibition were simulated to model the presence of IPSCs generated at either F1 or F2 synapses, both modelled with the exponential difference model, as shown in [1], but with IPSC_F1-fast_ (*τ_up_* lil 1.5, 2.25, 3.75 ms, *τ_down_* lil 4.0, 9.0, 12.0 ms, *σ_max_* = 0.25) or IPSC_F2-slow_ (*τ_up_* ∈ 1.2, 2.0, 2.5 ms, *τ_down_* = 20ms, *σ_max_* = 0.25). Delivery of the IPSC_F1-fast_ waveforms were delayed by 2 ms relative to the timing of the RGC inputs whereas the IPSC_F2-slow_ waveforms were delayed by 15 ms. Due to the skew in the underlying distributions, these latencies were based upon the median values that were obtained in the control and TTX optogenetic experiments as illustrated in Figure 2. Synaptic dynamics of IPSC_F1-fast_ was controlled with a function inversely proportional to the instantaneous frequency of the RGC stimulus pattern whereas the magnitude of the smaller IPSC_F2-slow_ inputs was maintained at all frequencies. All inhibitory drives were controlled by a non-linear leak conductance derived from the GHK equation to mimic the presence of an asymmetrical Cl^-^ ion distribution across the membrane with an internal concentration *C_in_* of 4 mM and an external concentration *C_out_* of 110 mM.

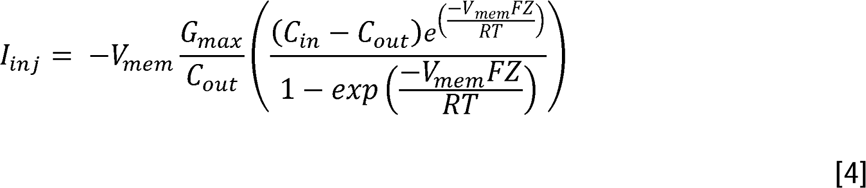

**Figure 1.**
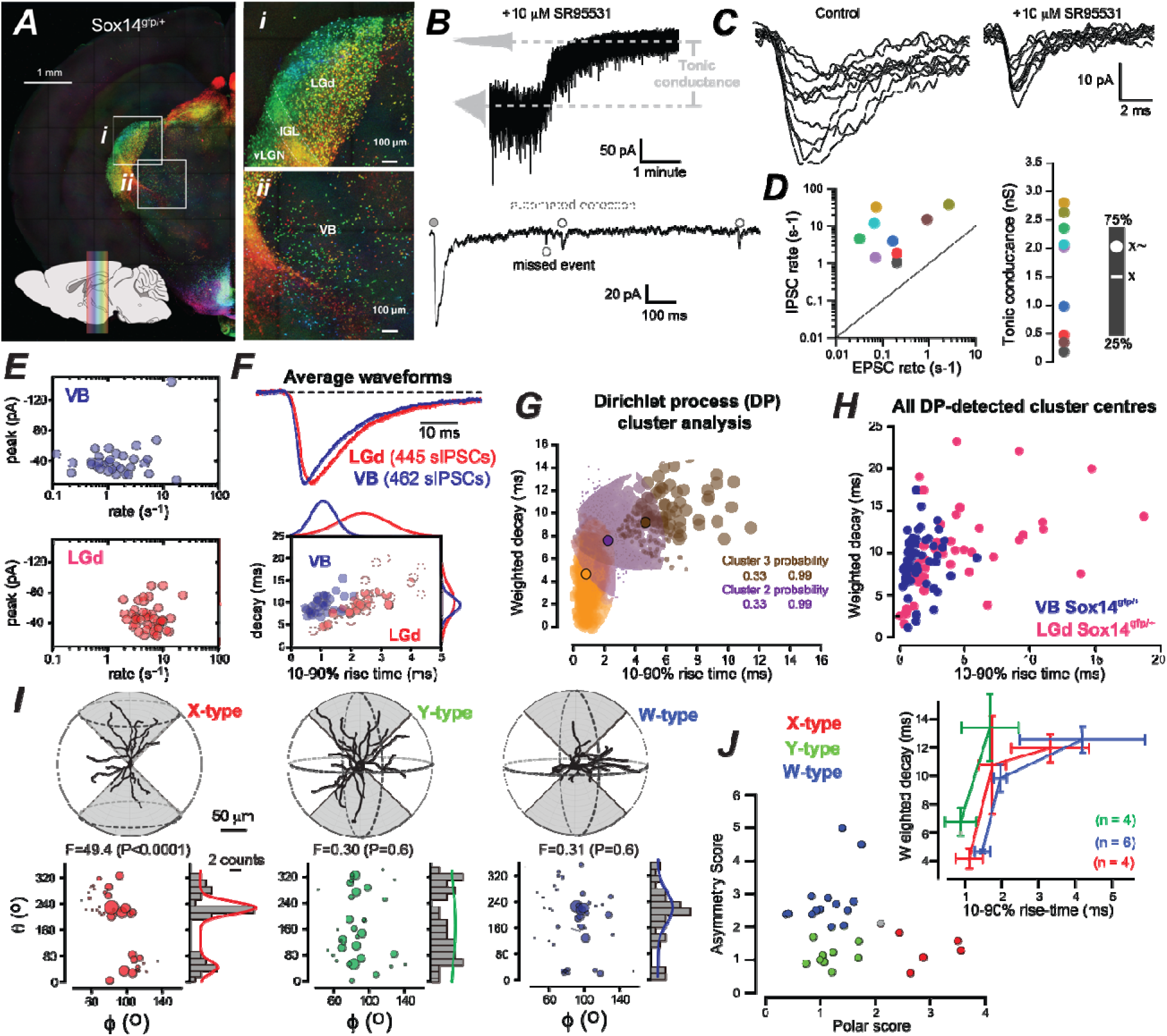
Slow-rising IPSCs are more prominent in TC neurons of the LGd. ***A***, Full projection of two-photon imaging data obtained following automated block face imaging of the Sox14^gfp/+^ mouse brain. Each fluorescently labelled Sox14^gfp/+^ cell is colour-coded according to rostro-caudal location in accordance with the colour sequence key superimposed on the sagittal brain section in the bottom left-hand corner of the panel. Higher magnification images of the dataset concentrating on the dorsal lateral geniculate nucleus (LGd) region and the ventrobasal (VB) region of the thalamus are shown in panel i and ii. Note the high density of Sox14^gfp/+^ cells in the intergeniculate leaflet (IGL) and ventral portion of the lateral geniculate nucleus (LGv) and the much lower density of Sox14 cells in the adjacent VB. ***B***, A continues current trace obtained during a whole cell voltage-clamp recording showing GABA_A_ receptor blockade in the presence of 10 μM SR95531. All point histograms (grey) were used to calculate the average holding current before and after GABA_A_ receptor blockade. The change in holding current was subsequently used to estimate the tonic conductance in these cells. The initial brief section of the current shown below the full trace illustrates the location of transient events detected with our automated template fitting routine (grey circles). The single open circle illustrates a missed transient that was much faster than the template. ***C***, Examples of transient events recorded in control conditions and during GABA_A_ receptor blockade. Note the much faster kinetics of the transient events recorded in the presence of 10 μM SR95531. ***D***, scatter plot of the sEPSC rates estimated in 9 cells during GABA_A_ receptor blockade versus the putative sIPSC rates estimated from these same cells in control conditions. Note how the sIPSC rates are consistently greater than sEPSC rates in each cell. The plot on the right shows the tonic conductance estimates obtained from these same 9 cells (colour coded) with a violin plot in grey and a bar graph showing the 25% and 75% range and the median (white circle) and mean (white line) values. ***E***, The scatter plots show the average sIPSC rate and average peak amplitude for sIPSCs calculated for 43 LGd relay neurons (red circles) and 32 VB relay neurons (blue circles) recorded in the presence of 10 mM kynurenate to block sEPSCs of the type shown in panel C.. Note the use of a log scale for plotting sIPSC rates. ***F***, Average sIPSC waveforms constructed from 445 sIPSCS recorded from an LGd relay neuron (red) compared to 462 sIPSCS recorded from a VB relay neuron (blue). The automated template fitting routine aligned events on the initial rising phase of the waveform as detected by a first derivative analysis. Note the faster monotonic rising phase of the VB waveform. The scatter plot below these traces indicates results from the entire population with 43 LGd cells and 32 VB cells with histograms aligned on the axis with Gaussian fits to illustrate the underlying distributions for rise-time and decay estimates. ***G***, Illustration of the variational Dirichlet process (DP) gaussian mixture model that was used to perform cluster analysis. The bubble plot demonstrates the probability that any point is associated with the 3 identified clusters. ***H***, A scatter plot of the centroid estimate for all clusters identified following DP-analysis for TC neurons in the LGd and VB of Sox14^gfp/+^ mice. ***I***, Scatter plot of morphological differences detected for thalamic relay neurons in the LGd. ***I***, Examples of three neurobiotin fills along with a graphical representation of the three different relay neuron morphologies. Dendrite branch points were quantified using a three-dimensional analysis of the x,y,z co-ordinates with the center of the soma set as 0,0,0 co-ordinates. The plot of I] and I] angles demonstrates the presence of two clear clusters for X-type relay neuron, consistent with a bi-polar distribution. The size of each circle denotes the distance from the soma. ***J***, Scatter plots comparing the the three morphological classes identified in the LGd along with an analysis of the weighted decay and 10-90% rise-time estimates for a subset of cells that had been classified based upon their morphology.

**Figure 2.**
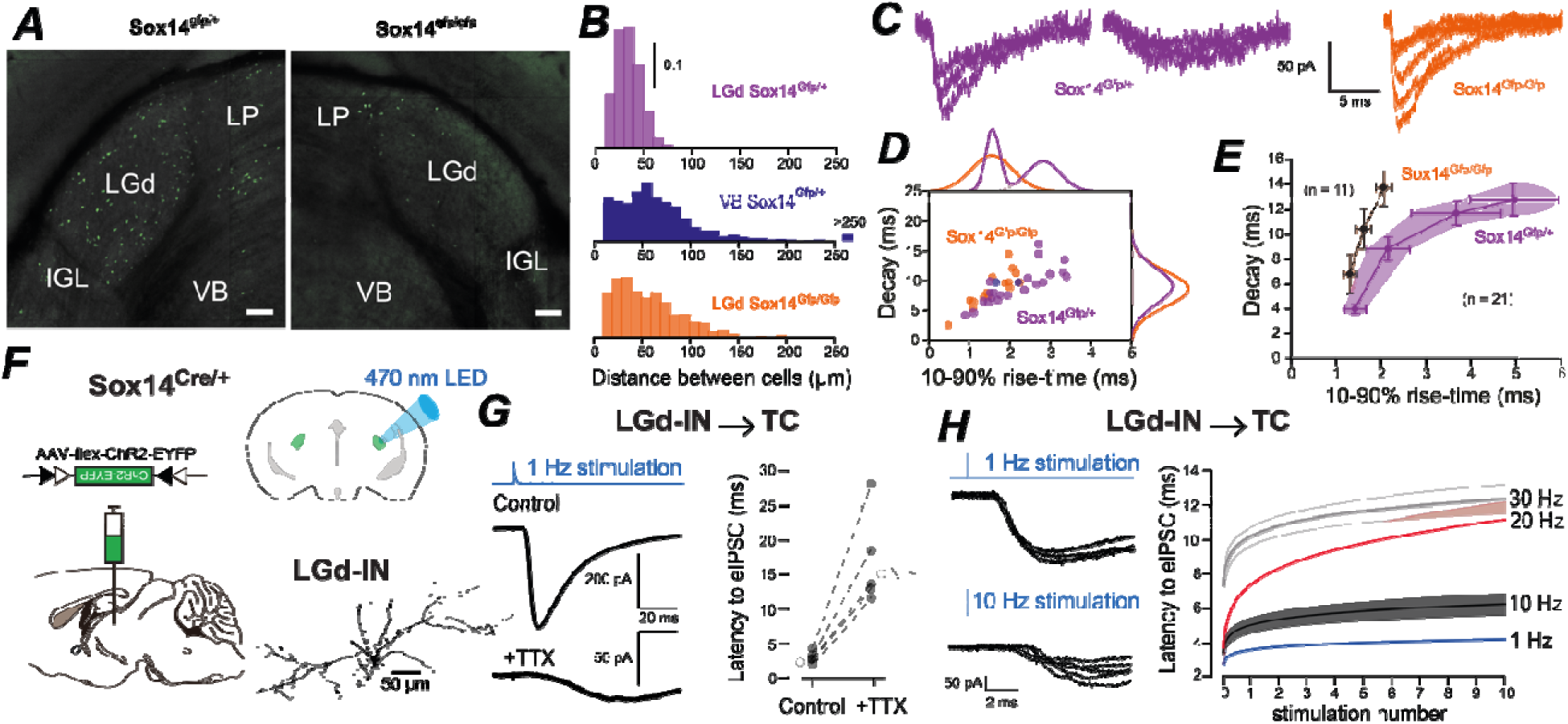
Interneurons are responsible for the slow-rising IPSCs onto TC neurons. . ***A***, Images showing qualitative comparison of Sox14^gfp/+^ and Sox14^gfp/gfp^ brain sections at AP −2.4. Sox14^gfp/gfp^ (a geneti knockdown) shows decreased GFP expression across midbrain, thalamic and hypothalamic regions. Labels indicate ventrobasal complex (VB), lateral posterior nucleus (LP), dorsal lateral geniculate nucleus (LGd), and the intergeniculate leaflet (IGL). Scale bars are 100 μm in length **B**, Quantification of nearest neighbour distances for Sox14 cells in the LGd and VB regions of the thalamus and comparison to the LGd region in Sox14^gfpgfp^ mice. The histograms are probability density functions for the nearest neighbour distances calculated for the LGd (purple) and VB (blue) regions of the Sox14^gfp/+^ brain with bin widths of 10 μm reflecting the minimum sampling resolution used for these imaging experiments. The bottom histogram is the quantification of Sox14 neurons in the LGd of Sox14^gfpgfp^ mice (orange). ***C***, Examples of detected sIPSCs recorded from TC relay neurons in the LGd of Sox14^Gfp/+^ (purple) and Sox14^Gfp/Gfp^ (orange) mice recorded at a command voltage of −70mV. ***D***, Scatter plot comparing 10-90% rise-time and weighted decay between Sox14^Gfp/+^ (purple) and Sox14^Gfp/Gfp^ (orange) LGd relay neuron population. Each dot represents averaged 10-90% rise time and weighted decay values for a single recorded neuron. Note that the Sox14^Gfp/+^ population has significantly slower 10-90% rise time compared to Sox14^Gfp/Gfp^ neurons, while there is no significant difference in weighted decay. ***E***, Data from cluster analysis. The scatter plots show the average values for each cluster center for all data gathered from the two strains. Note the absence of slow-rising clusters in the Sox14^Gfp/Gfp^ (orange) data. **F**, Illustration of the strategy used to deliver ChR2 into LGd-INs in the Sox14^Cre/+^ mouse line and optical stimulation in the acute slice preparation. The 470 nm LED output (blue trace) recorded from the water immersion objective lens that was used to excite ChR2-expressing LGd-INs. The morphology of a single LGD-IN recorded from the Sox14^Cre/+^ line is also shown. ***G***, Example of current traces recorded from a TC neuron responding to LGd-INs excitation following brief light pulses at 1 Hz in control condition and in the presence of 1 μM TTX. Note the slower kinetics and onset of the eIPSC recorded in the presence of TTX. This is consistent with the reported properties of dendro-dendritic GABA release. The scatter plot shows the **l**atency of eIPSCs to brief 1Hz light pulse stimulation in control condition and in the presence of TTX in the same neurons. The open circles denote the median values obtained for these skewed distributions. ***H***, Example traces recorded from the same TC neuron at the end of the 1 Hz and 10 Hz stimulation protocols recorded in control conditions. Note the long latency of the eIPSCs recorded at the higher stimulation rate. The right-hand plot shows the change in eIPSC latency during LED stimulation rates of 1, 10, 20 and 30 Hz. The solid line is the average fit to data from five individual recordings and the shaded areas are the SEM of these fits.

Where *G*_*max*_ is the maximum conductance of the leak, *R* is the gas constant, *F* is the Faraday constant, *T* is absolute temperature and *Z* is the valence of the ion.

### Data analysis

#### Voltage-clamp experiments

Spontaneous IPSCs were detected using a scaled template matching procedure implemented in WINEDR (Version 3.0.9) kindly provided by John Dempster (John Dempster; University of Strathclyde, UK) The template matching routine was optimized for the detection of sIPSCs using a template with 1 ms rising phase and a 10 ms decay phase. Waveform averages were constructed from sIPSCs that exhibited monotonic rises and uninterrupted decay phase. The detected events used to construct the average waveform were aligned on the initial rising phase using a first derivative to identify the initial current deflection. Average baseline current levels were calculated during a 10 ms epoch immediately before each detected event and the peak amplitude was determined relative to this value. The decay of individual sIPSCs was calculated as the charge transfer during the baseline corrected sIPSC divided by the sIPSC peak amplitude. Cluster analysis for the 10-90% rise-time and weighted decay distributions was undertaken in MATLAB (MATLAB, 2019).

A variational Dirichlet process (DP) gaussian mixture model was used to enable automatic determination of an appropriate number of mixture components with nonparametric Bayesian techniques. (Sato et al., 2012; Kurihara 2008). For DP cluster analysis a minimum of 100 automatically detected sIPSCs were needed in each recording. We decided on a non-parametric Bayesian model for clustering that allowed for flexible and data-driven clustering without making assumptions about the number of clusters or their shapes. Unlike traditional clustering algorithms that assume a fixed number of clusters or rely on predefined parametric models (e.g., Gaussian Mixture Models), non-parametric Bayesian techniques allow the number of clusters to be determined from the data itself. The underlying algorithm uses the Dirichlet process which is a fundamental concept in non-parametric Bayesian clustering. This technique allowed us to separate the 10-90% rise times and weighted decay distributions into clusters without predefining the number of clusters beforehand as shown in Figures 1D and 1E. Output from the software gives the number of clusters found, together with the centroids of each component cluster and its associated covariance matrix.

#### Current-clamp experiments

The input-output relationship of TC neurons was determined by varying g_max_ during the delivery of the EPSC_AMPA_ and EPSC_NMDA_ waveforms to simulate variability in failure rate at the retinogeniculate synapse. The timing of APs in each TC neuron was detected by threshold crossing at 0 mV and the resulting input-output relationship was used to estimate the excitatory conductance required to produce 50% of the maximum number of APs in the relay neuron (*EXC_50_*). The influence of the 3-types of inhibition, delivered singularly or in combination, were determined on the input-output characteristics of relay neurons during delivery of the *EXC_50_*.

All input-output relationships were fitted with a descriptive Boltzmann function of the form:

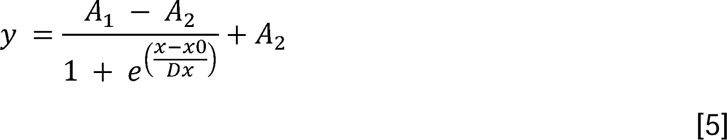

Where *A*_1_ is the initial value of the fit, *A*_2_ is the final value, *x*_0_ is the centre of the fit and *Dx* is the slope constant.

#### Latency to First Spike

Input clusters were objectively defined for each RGC pattern as one or more inputs separated by a time interval smaller than the mean inter-event interval in a similar manner previously described (Mazzoni *et al*., 2007; Xu & Baker, 2018). The neuronal latency to first spike (LTFS) for each input cluster was calculated as the time difference between the first input of the input cluster and the first output within the defined cluster. The LTFS was calculated at increasing synaptic weights to elucidate the relationship between LTFS and input conductance.

#### Rate Coding

Rate coding characteristics of the neuron was calculated by dividing the total number of outputs for each burst of inputs by the duration of the respective burst. This was calculated for each burst at increasing levels of conductance magnitudes to determine the filtering properties of the neuron and the effects of inhibition on this parameter of neuronal information processing.

#### Information Loss

The input-output relationship for the whole RGC trace was determined by varying g_max_ and calculating the timing of the evoked action potentials with threshold crossing at 0 mV. A simple analysis of residuals approach was used to estimate the proportion of the variance in the AP intervals that was predicted from the variability in the input. The inter-event intervals for the RGC input pattern were determined at a 5 ms bin width and the AP inter-event interval distribution at each *g_max_* was used to calculate the goodness of fit (R^2^) using the sum of squares to calculate the unexplained variance (SSres) and the total variance (SStot) according at each *g_max_* was calculated according to *R^2^ = 1-(SSres/SStot).* Information Loss (IL) is then defined as the remaining information that was not transferred, hence the formula: *IL = 1 – R^2^*.

TC neurons alternate between tonic and burst firing modes depending on the activity of Ca^2+^-dependent channels located in the membranes of the soma and dendrites (Llinas & Jahnsen, 1982). In tonic firing mode, the transfer of information arriving from EPSCs is generally linearly associated with the output of the TC neuron, however in burst firing mode, the relationship between the input and output is non-linear as the number of outputs exceed the number of inputs (Sherman, 2001). Thus, we excluded TC neurons that were observed to switch to burst firing mode; as evidenced by AP numbers greater than the number of RGC inputs.

#### Neuronal morphology

Following electrophysiological recording, slices were fixed in 4% PFA, permeabilized with 0.2% Triton-X at room temperature for 1-2 hours and non-specific binding was blocked with 5% donkey serum in PBS. A 1:200 dilution of streptavidin-Alexa555 (Invitrogen^®^, USA) was then conjugated to the biocytin and slices were washed in PBS and mounted in Vectashield^®^ H-1000 (refractive index = 1.44). Individual neurons were imaged using a Zeiss LSM 510 CLSM microscope (Facility for Imaging by Light Microscopy, FILM, Imperial College), with a He-Ne 543nm laser and filter settings optimised for an Alexa-555 emission peak at 565 nm. The numerical aperture of the 40x oil-immersion objective was 1.3, the emission wavelength was 700 nm, and the refractive index was of our mounting medium was 1.516. Given these parameters, the optical section thickness was set at 1.02 μm. The confocal pinhole was set to one Airy unit and scan speeds were typically set to 9.3 seconds/slice with the photomultiplier gain adjusted to avoid saturation of the fluorescent signal. Optical sections were over-sampled in the *x, y* and *z* dimensions (0.1 μm, 0.1 μm, 0.49 μm) to satisfy the Nyquist theorem and each pixel intensity was recorded at 32-bit resolution to optimize dynamic range.

Optical sections obtained following confocal imaging were imported into Reconstruct^®^ (Fiala, 2005) where the *x, y* and *z* Cartesian coordinates of each dendrite branch point and end point were normalized with respect to the soma centre and converted into spherical coordinates according to 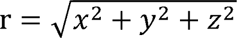 and θ = arctan 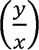, and = arccos 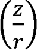. A *k*-means cluster analysis was employed to negate the mpact of cell *φ* orientation that could potentially influence the outcome of more conventional Sholl analyses (Friedlander *et al*., 1981). The radial distribution of dendrites in Y-type thalamic relay neurons resulted in a low F value for the θ and φ angle distributions with no statistical evidence of two clusters. In contrast, the polar distribution of dendrites that characterizes X-type thalamic relay neurons resulted in two clusters in the θ and φ angle distributions with a high F value that was considered significant at P<0.05 following analysis of variance (ANOVA). The size of the bubble in the scatter plots constructed from this analysis indicated the radial distance calculated for each point in the θ and φ angle distributions. The θ and φ angles for the data in each cluster could also be used to calculate the average distance between the two clusters. The asymmetry score was calculated from the ratio of counts in the highest and lowest density hemispheres with a value greater than 2 being considered asymmetrical. As shown in Figure 1H, TC relay neurons were characterized into X, Y and W cell types based upon the results of the polar and asymmetry scores. X-type relay neurons were symmetrical (Asymmetry score <2) with the presence of two distinct clusters (Polar score >2). Y type relay neurons were symmetrical (Asymmetry score <2) but non-polar (Polar score <2). W-type relay neurons were asymmetrical (Asymmetry score >2) and non-polar (Polar score <2).

#### Statistical tests

All average values represent the mean ± the standard error of the mean (SEM). Data distributions were compared using Origin 8.5 (Microcal, Century City, CA) and functions were fitted to data distributions using unconstrained least-squared fitting procedures. A *k*-means cluster analysis was also performed on the θ and φ angle distributions calculated from the *x, y* and *z* Cartesian coordinates of dendrite branch points and end points. Two significant clusters were considered to be present in the distribution when the F value associated with the analysis of variance (ANOVA) reached a P<0.05. The custom code used can be found in the following GitHub directory: https://github.com/dd119-ic/in-dlgn-paper.

## Results

We have previously reported that slow-rising sIPSCs are mediated by γ1 subunit-containing GABA_A_ receptors expressed at the dendro-dendritic F2-type synapses formed by local LGd interneurons (Ye *et al*., 2017). This suggestion was consistent with previous data demonstrating that calcium spikes in the dendrites of LGd interneurons result in delayed asynchronous GABA release at these dendro-dendritic synapses (Acuna-Goycolea *et al*., 2008). Given these observations, we predicted slow rising IPSCs would be less evident in the ventrobasal thalamus (VB) where interneuron density is known to be lower (see **Figure 1A**). Before quantifying sIPSC properties we have included a description of experiments undertaken without glutamate receptor blockers in the recording solution to illustrate the properties of sEPSCs recorded during whole-cell voltage-clamp recording at a command voltage of −70mV. The GABA_A_ receptor block was undertaken to isolate sEPSCs but also enabled the tonic conductance to be measured and compared to previous published data. The holding current was measured in control conditions and in the presence of 10 μM SR95531. At room temperature, the tonic conductance was 1.5 ± 0.3 nS (*n*=9) with a similar range of values to those reported by our laboratory at physiological temperatures (Bright *et al*., 2011). This observation is intriguing given that we and others have shown that faster rates of deactivation/desensitization are an obvious feature of high temperature recording (Houston *et al*., 2009; Bright *et al*., 2011). The bottom trace shown in **Figure 1B** illustrates how our automated detection protocol often failed to detect the very fast current transients that were present in control conditions. As shown in **Figure 1C**, these fast EPSCs that were recorded in the presence of 10 μM SR95531 occurred at a low rate of 0.48 ± 0.28 s^-1^ (*n*=9 cells). This compared to a rate of 12.0 ± 4.5 s^-1^ for putative sIPSCs that were automatically detected in the same cells (see **Figure 1D**). In these nine cells, the sEPSCs were characterized by a 10-90% rise-time of 0.97 ± 0.16 ms, a peak amplitude of −26.2 ± 4.9 pA and a weighted decay of 5.8 ± 0.9 ms. In all subsequent voltage-clamp experiments the detection of sIPSCs was performed in the presence of the broad-spectrum glutamate receptor antagonist kynurenate (10mM).

### Slow-rising IPSCs are a feature of X- and W-type relay neurons of the LGd

Whole-cell voltage-clamp recordings made at a command voltage of −70mV in the presence of broad-spectrum glutamate receptor blockers were obtained from 43 TC neurons in LGd and 32 TC neurons in VB (**Figure 1E**). The kinetic properties of automatically detected sIPSCs were analyzed in each cell with an average of 460 ± 64 sIPSCs detected in TC neurons from LGd and 403 ± 66 sIPSCs in VB. There was little difference in the average peak amplitude of sIPSC recorded in the two thalamic regions (−47.6 ± 2.6 pA in LGd versus −40.6 ± 4.0 pA in VB; t-Test, *P* = 0.14). The mean instantaneous rate of sIPSCs in the LGd was 6.5 ± 0.6 s^-1^ compared to 2.7 ± 0.7 s^-1^ in VB (t-Test, *P* = 0.0001). From the scatter plots shown in Figure 1E, the main difference between LGd and VB recordings was the greater variability in sIPSC rates observed in VB and an absence of cells with rates under 1 s^-1^ in the LGd dataset. Also note how VB recordings did contain cells with high sIPSC rates of >10 s-1. Overall, the 10-90% rise-time of the sIPSC waveforms (**Figure 1C**) constructed from 43 dLGN relay neurons was 2.42 ± 0.13 ms compared to 1.08 ± 0.07 ms for 32 relay neurons recorded from VB (t-Test, *P* < 0.005). However, the average sIPSC weighted decay was not significantly different (t-Test, *P* = 0.49) between LGd (9.93 ± 0.66 ms) and VB (9.44 ± 0.3 ms). Scatter plots of the 10-90% rise-time and weighted decay of all sIPSC in each cell was constructed and a variational Dirichlet process (DP) gaussian mixture model used to estimate the centroids of each component cluster (**Figure 1D**). Application of this non-parametric Bayesian technique to mixture modelling avoids *a priori* decisions on the number of clusters. The average number of distinct cluster types detected from DP analysis was 2.4 ± 0.2 for LGd recordings (21 cells) compared to 1.8 ± 0.5 in VB (30 cells) demonstrating a significant reduction in the number of cluster types (un-paired t-Test p=0.015 with Welch correction for unequal variance). However, when all detected clusters were pooled (51 clusters in LGd form 21 cells compared to 55 clusters in VB from 30 cells) we can see also see that the 10-90% rise times was significantly slower (un-paired t-Test p=8.8E-5 with Welch correction) in the LGd (4.1 ± 0.6 ms) compared to VB (1.6 ± 0.2 ms). However, as expected from the previous analysis of the kinetics of average waveforms, the weighted decay of the DP detected clusters was not significantly different (un-paired t-Test p=0.98 with Welch correction) between LGd (9.61 ± 0.7 ms) and VB (9.63 ± 0.3 ms). However, it should be noted that the similar decay kinetics of sIPSCs recorded from TC neurons in LGd and VB (**Figure 1E**) is contrary to a previous report (Yang *et al.,* 2017). The reason for this discrepancy is not clear but the DP analysis provides an unbiased identification of heterogeneity within the sIPSC population. Based upon polar and asymmetry scores we classified 29 TC neurons within the LGd into 5 X-type, 10 Y-type and 14 W-type TC neurons (**Figure 1F**). X-type TC neurons exhibited bipolar dendrite distributions, with most dendrite branch points residing in one axis. Y-type TC neurons had radial symmetry of dendrites and no orientation preference. W-type TC neurons had most of their dendritic tree contained within one hemisphere with clear asymmetry and low polarity (**Figure 1G**). When this quantitative morphological analysis was combined with DP-based sIPSC cluster analysis, it became clear that slow-rising IPSCs are a feature of both X and W-type TC neurons but are absent from Y-type TC neurons of the LGd (**Figure 1H**).

### Local interneurons contribute to slow-rising IPSCs

We have previously shown that reducing LGd interneuron numbers in the loss-of-function Sox14^gfp/gfp^ mice was associated with a reduction in sIPSC rates (Jager *et al.,* 2016). Automated cell counting following serial 2P imaging (Jager *et al*., 2021) was used to compare the Euclidian distance between interneurons in Sox14^gfp/+^ and Sox14^gfp/gfp^ mice (**Figure 2A**). Nearest neighbor plots constructed from these 3D measurements (**Figure 2A**) indicated a maximum probability of 25 μm between interneurons in the LGd compared to 55 μm in the VB of Sox14^gfp/+^ brains. No Sox14^gfp/+^ cells were found further than 85 μm apart in the LGd compared to separations of >205 μm in VB (**Figure 2B**). The distribution of Sox14 cells in the brains of Sox14^gfp/gfp^ mice exhibited a distribution that was more like the distribution observed in the VB of the Sox14^gfp/+^ brains (**Figure 2B**). The sIPSCs we record from TC neurons can result from GABA release from reticular axons as well as local interneurons. At present, we propose that the reduction in LGd interneuron density in Sox14^gfp/gfp^ mice is responsible or the reduction in the prevalence of slow-rising sIPSCs. However, we cannot rule out the possibility that some form of developmental compensation has taken place in response to reducing local interneuron density. Whole-cell recordings from 15 TC neurons in the LGd of Sox14^gfp/gfp^ mice were compared with data from 21 littermate Sox14^gfp/+^ controls. Following automatic detection of sIPSCs there was a conspicuous lack of slow rising sIPSCs in Sox14^gfp/gfp^ TC neurons recorded from the LGd making this population qualitatively similar to data obtained from the VB (**Figure 2C**). Quantitatively, the 10-90% rise-time in control Sox14^gfp/+^ TC cells was significantly slower (2.16 ± 0.17 ms, n = 21) compared to data from Sox14^gfp/gfp^ recordings (1.57 ± 0.11 ms, n = 15, t-Test, *P* <0.05). In contrast, the average IPSC weighted decay estimates were not significantly different (9.11 ± 0.68 ms in control versus 8.63 ± 0.65 ms in the Sox14^gfp/gfp^ recordings, t-Test, *P* = 0.65). The distribution of the average rise-time data collected in the control group was best described with the sum of two Gaussians suggesting two populations of relay neurons (**Figure 2D**). In the Sox14^gfp/gfp^ strain the distribution of rise-times were well described by a single Gaussian fit with a peak of 1.6 ms consistent with a lack of slow-rising sIPSCs. In contrast, the distribution of decay times was similar in the two strains with a single Gaussian fit of 8.9 ms and 8.6 ms. Cluster analysis was once again applied to the sIPSC data recorded from individual relay neurons. In the littermate control strain (Sox14^gfp/+^), all IPSC distributions were like those observed in the LGd of the previously analyzed C57Bl6 strain. Four discreet IPSC clusters were detected in each distribution (*n* = 21) with clear evidence of a slow-rising sIPSC population (**Figure 2E)**. However, the knockdown mice (Sox14^gfp/gfp^) most cells displayed only three clusters with an absence of the slow-rising sIPSC population, reminiscent of the relay neurons in VB thalamus (**Figure 2E**).

### Optogenetic activation of GABA release from dLGN-INs generates fast and slow IPSCs

Virally delivering Cre-dependent ChR2-YFP constructs into the LGd of the Sox14-Cre mouse enabled the optogenetic stimulation of GABA release from LGd interneurons that resulted in a postsynaptic response in adjacent TC neurons (**Figure 2F**). As expected (Jager *et al*., 2016), a ChR-2 mediated conductance was present in LGd interneurons recorded 2-3 weeks after transfection (**data not shown**) and this excitatory conductance was sufficient to elicit robust non-accommodating APs at stimulation frequencies of up to 20 Hz. While AP-induced depolarization can propagate globally to promote GABA release throughout the interneuron dendrites and axons, pharmacological evidence suggests that local dendritic GABA release is mediated through activation of L-type calcium channels in an AP-independent-manner (Acuna-Goycolea *et al*., 2008; Casale & McCormick, 2011; Crandall & Cox, 2012). By blocking AP-dependent GABA release from interneurons with 1 μM TTX, AP-independent eIPSCs with slow onset and rise-time were unmasked in 5 out of 7 whole-cell voltage-clamp recordings at −70mV (**Figure 2G**). The remaining 2 recordings did not show any eIPSCs in the presence of TTX (data not shown). At LED stimulation rate of 1 Hz the average latency of eIPSCs was 3.28 ± 0.39 ms (*n*=5) in control condition. In the presence of 1 μM TTX, the average latency of unmasked dendrodendritic eIPSCs was 16.95 ± 3.03 ms (**Figure 2G**). After characterizing interneuron mediated fast and slow IPSCs, we decided to look at the temporal pattern of phasic IPSCs evoked by brief light pulses at different frequencies. We have previously reported the tonic current at 10, 20 Hz stimulus frequency is mediated by extrasynaptic δ-containing GABA_A_Rs (Jager *et al*., 2016). Here, we report a gradual increase of eIPSC latency at different stimulus frequencies over 10 seconds of stimulation period (**Figure 2H**). The average latency of eIPSCs did not change significantly at 1Hz stimulation rate. However, at LED stimulation rates of 20 Hz, the average latency increased from 3.4 ± 0.1 ms to 10.3 ± 0.8 ms (n = 5) during the 10 second stimulation period. The slow eIPSCs towards the end of 20 Hz stimulation was very similar to the slow eIPSCs recorded in the presence of TTX, suggesting fast eIPSCs originating from axons were fast depressing, while the slow dendritic eIPSCs remained at higher frequencies.

### TC neurons respond to different RGC patterns with similar input sensitivity while preserving AP output differences

Dynamic-clamp (**Figure 3A**) offers the ability to simulate the presence of excitatory and inhibitory synaptic conductance changes in ways that are not possible with other techniques. For example, the conductance change associated with optogenetic activation of ChR2 involves the opening of non-specific cation channels and, therefore, the I/V relationship of membrane permeability changes cannot be altered during an experiment. Using the dynamic-clamp approach we can vary the underlying I/V relationship of the conductance change and specify the exact timing and underlying kinetics of this conductance change. Therefore, we can introduce precisely timed levels of excitation and inhibition to TC neurons to simulate the presence of visually driven feedforward inhibition and compare the influence of F1- and F2-type synapses on the gain of TC neurons as well as introduce a conductance to mimic the tonic conductance known to present on these TC neurons (**Figure 3A**).

**Figure 3.**
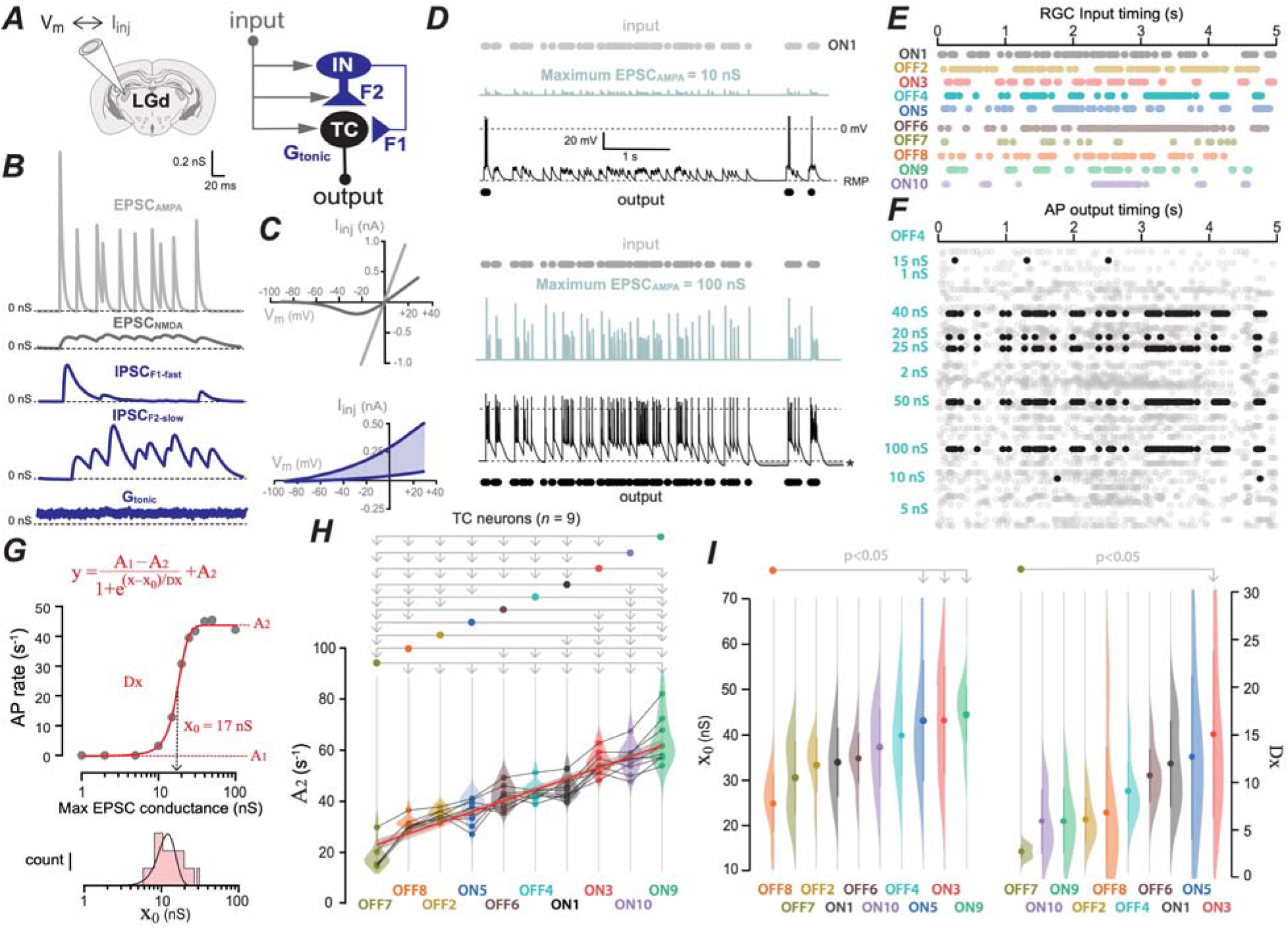
TC neurons can distinguish between different RGC input patterns. ***A***, Illustration of simultaneous voltage monitoring and current injection used to mimic the conductance changes associated with the thalamic circuitry. The retinogeniculate input pattern was used to control the conductance waveforms generated by the feedforward circuit associated with local interneurons at both axo-somatic F1 and dendro-dendritic F2 synapses. The final conductance waveform was designed to simulate the tonic conductance present on TC neurons. ***B***, Time-varying conductance waveforms used to mimic the AMPA- and NMDA-type retinogeniculate synapses (EPSC_AMPA_ and EPSC_NMDA_) are shown in the top two gray traces, whereas the axo-somatic F1-type fast synapses (IPSC_F1-fast_), the dendro-dendritic F2-type slow synapses (IPSC_F2-slow_) and the persistent tonic conductance changes (G_tonic_) are shown in blue. ***C***, The upper I-V relationships were used to calculate the current injection at any given membrane voltage for the linear Ohmic relationship used to control the AMPA-type EPSC waveforms compared to the more complicated Boltzmann function used to control the non-linear voltage-dependence of the NMDA-type EPSCs shown in B. The lower blue traces are the I-V plots of the GHK-type open rectification used to simulate the chloride ion flux through GABA_A_ receptors. The shaded blue region depicts the range of I-V relationships used to control the IPSC_F1-fast_, IPSC_F2-slow_ and G_tonic_ time varying conductance changes. ***D***, Illustration of a dynamic-clamp experiment delivering an ON-type RGC input pattern (labelled ON1 in panel E) at a maximum conductance of 10 nS. The timing of the input pattern is shown with filled grey circles and the resulting time-varying conductance change used to mimic the retinogeniculate synapse onto this TC neuron is shown in with the light blue trace. The black trace is the membrane voltage recorded from the TC neuron during this stimulation pattern and the timing of each AP is shown with black circles below each trace. In this neuron, only 7 APs were elicited when the maximum excitatory conductance was delivered at 10 nS. ***E***, The same cell shown in ***D***, but the maximum excitatory conductance has been raised to 100 nS. An AP is now elicited in response to every EPSC. Note also how a prolonged membrane hyperpolarization was observed once this maximum excitation was removed (asterisk) that we assume reflects a Ca^2+^ activated potassium conductance triggered by sustained high frequency AP firing. ***F***, Raster plot of the timings used to simulate the 10 different RGC input patterns used in subsequent experiments. Each input pattern is colour coded. ***G***, Raster plots illustrating all AP timings recorded from a single relay neuron in response to 5 second dynamic clamp protocols simulating 10 separate RGC timing patterns delivered at varying maximum conductance values. The solid black circles mark the timing of this neuron to a single pattern of RGC inputs named OFF4 at maximum conductance values ranging from 1 to 100 nS. ***H***, Plot of the input-output relationship constructed from the AP firing of a single relay neuron in response to the OFF4 RGC input pattern. This data was fitted with the Boltzmann function shown in red from which the maximum AP firing rate (A_2_), the inflection point of the curve (x_0_) and the slope coefficient (Dx) was extracted. Below the I-O plot is a histogram showing the distribution of x_0_ values obtained for 15 relay neurons stimulated with the same ON1 type input pattern. ***I***, The A_2_ values obtained from 9 TC neurons that received all 10 RGC input patterns are plotted as scatter and violin plots. The solid lines connect data obtained from individual relay neurons. Data in this plot were ranked according to the mean A_2_ values and ANOVA followed by Bonferroni correction was used to determine significance between response types. The red line is the result of a linear regression with a Pearson’s r of −0.9 indicating the ability of this ranking to explain much of the variability in the maximum firing rates. The shaded area around the regression line is the 95% confidence limits for this fit. ***J***, Comparison of x_0_ and Dx distributions obtained for all 10 RGC input patterns delivered to the same 9 TC neurons in I. The RGC ranking was sorted based on either the mean x_0_ or mean Dx values and ANOVA demonstrated how few RGC patterns were significantly different from each other when considering either parameter. For example, based upon Dx values, only the TC neurons sensitivity to OFF7 and ON3 were significantly different. Linear regression analysis further demonstrated a lack of correlation between this ranking and the variability in either x_0_ or Dx.

The timing of EPSCs used to mimic retinogeniculate input onto TC neurons was based upon extracellular recordings of RGC firing patterns generated in response to a natural scene projected onto a mouse retinal preparation (Meytlis *et al*., 2012). The kinetics of the EPSC_AMPA_ and EPSC_NMDA_ waveforms (**Figure 3B**) was based upon voltage-clamp data obtained at the retinogeniculate synapse (Blitz & Regehr, 2003). The peak amplitude of each EPSC was varied according to the properties of frequency-dependent depression previously described at the retinogeniculate synapse (Chen *et al.,* 2002). The AMPA to NMDA ratio was set such that each NMDA peak amplitude was 10% of the associated AMPA peak amplitude. During delivery of the EPSC_AMPA_ waveform, the dynamic-clamp software calculated the injected current at a given voltage according to a simple Ohmic leak relationship, whereas a Boltzmann function was used to mimic the non-linear behavior of NMDA channels (**see Figure 3C**). Once threshold was reached, injection of the summated current calculated from the combined EPSC_AMPA_ and EPSC_NMDA_ waveforms resulted in robust AP firing in all relay neurons examined. For the example shown in Figure 3, only 7 APs were elicited in response to the excitation dynamic-clamp protocol that was based upon the ON1 RGC firing pattern that contained 109 retinogeniculate EPSCs. In the example recording shown in **Figure 3D**, the maximum EPSC conductance was only 10 nS. However, in **Figure 3E** we show the same cell when the maximum conductance was raised to 100 nS. In this situation, a total of 116 APs were elicited in response to the 109 EPSC inputs. The excess APs reflected burst firing at the end of this protocol due to the prolonged membrane hyperpolarization that was observed in this cell. This behavior is likely to reflect the presence of a Ca^2+^ activated potassium conductance triggered by sustained high frequency AP firing. We developed an experimental protocol that simulated-the arrival of 10 different RGC input patterns (**Figure 3F**) at varying synaptic weights from a maximum synaptic conductance of 1 nS to 100 nS. A total of 110 waveforms were randomly delivered to each TC neuron and the response to individual RGC patterns was extracted for subsequent analysis. For example, in the raster plot shown in **Figure 3G**, we have isolated the TC neurons response to the OFF4 input pattern. From this data, we constructed I-O relationships for each RGC pattern and examined the relationship between the synaptic weighting and the AP firing rate (**Figure 3H**). The resulting sigmoidal relationship was well described by a Boltzmann function from which we could extract the maximum AP firing (A_2_) for each TC neuron. The x_0_ and Dx values reflect the input sensitivity of each TC neuron to these RGC inputs. Specifically, the inflection point x_0_ indicates the synaptic weight that gives a 50% increase in AP firing rate while the slope coefficient Dx reports the change in synaptic weight that results in an e-fold change in AP firing rate. Therefore, the Boltzmann function was used to evaluate the output rate of each TC neuron to the 10 different RGC patterns as well as the input sensitivity of each TC neuron to these inputs.

The A_2_ distributions from each TC neuron was used to rank the RGC input patterns from the lowest to the highest output and linear regression demonstrated that this ranking explained 90% of the variability in maximum AP firing rates (**Figure 3I**). Not surprisingly, given that TC neurons encode visual information in the LGd, the AP firing rates in response to these RGC firing patterns were significantly different from one another; as evidenced by ANOVA followed by Bonferroni correction for repeated measures (**Figure 3I**). In contrast, if a similar ranking of TC neurons was undertaken based on either x_0_ or Dx we saw very little difference between RGC input patterns (**Figure 3J**). For example, based upon x_0_ values, only OFF8 was shown to be different to ON3,5 &9 and when considering Dx values, only a TC neurons sensitivity to OFF7 and ON3 were significantly different. Furthermore, linear regression analysis demonstrated a lack of correlation between the ranking and variability in either x_0_ or Dx. This result demonstrates that although the AP firing patterns of TC neurons robustly reflect differences in the timing of the RGC input patterns the input sensitivity of each TC neuron to these different RGC patterns is similar. This was an important consideration when interpreting subsequent data on the influence of inhibition on information transfer in response to different RGC patterns.

### Fast, slow and tonic inhibition are all capable of altering TC neuron excitability

The dynamic-clamp approach was used to simulate the presence of F1-type and F2-type synaptic inhibition as well as the tonic inhibition resulting from the continuous activation of extra synaptic GABA_A_ receptors. The current-voltage relationship chosen to calculate the injected current at the recorded membrane voltage was based upon the Goldman-Hodgkin-Katz (GHK) relationship to mimic the presence of a non-linear conductance of the type associated with the chloride ion flux through GABA_A_ receptors (**Figure 3C**). Based upon our voltage-clamp data, the slower rising and slower decaying F2-type IPSCs were generated with a 50% reduction in peak amplitude compared to the faster F1-type IPSCs. As shown in **Figure 3B**, the timing of each IPSC_F1-fast_ was delivered at the same time as the corresponding EPSC_AMPA_. This arrangement was chosen to simulate the presence of fast feedforward inhibition generated due to the retinogeniculate input onto LGd interneurons (Blitz & Regehr, 2005). Variability in the peak amplitude of IPSCs in the F1-type waveforms was calculated using the same relationship used to introduce peak amplitude variability into the EPSC_AMPA_ waveforms. Therefore, we are assuming that relay neurons and LGd interneurons share excitatory input from overlapping RGC receptive fields and that the retinogeniculate inputs to these two neuronal types have similar synaptic dynamics. In contrast, the unusual properties of dendro-dendritic GABA release from the slower F2-type inputs included a 5-10 ms delay before the injection of slow-rising and slow-decaying IPSCs with no reduction in the IPSC peak amplitudes at the different inter-event intervals. This behavior was based upon our observation on IPSC properties and the results of optogenetic stimulation of LGd interneurons (**Figure 2H**).

After determining x0 values for the excitatory drive (**Figure 3G**) we systematically varied the maximum conductance of the three different types of inhibitory conductance (**Figure 4A**) and then calculated the change in the average AP probability following delivery of all RGC input types (**Figure 4B**). From this analysis, it was apparent that introduction of F1-type inhibition was the least effective in reducing the average AP probability and tonic inhibition was the most potent. The dynamic-clamp approach further enabled the AP probability to be calculated at each inter-event interval (**Figure 4C**) to examine changes in the filtering characteristics of TC (**Figure 4D**). The density of the shaded bars in the resulting AP probability histograms (**Figure 4D**) indicate the number of events used to calculate the AP probability at each interval. This analysis demonstrates exactly how the low pass filtering characteristics of TC neurons was altered by the different types of inhibition. With excitation only the output IEI has a clear absence of events above 100 Hz even through these high frequency IEIs are present in the input histogram. This observation clearly illustrates the low-pass filtering characteristics of TC neurons. The introduction of F2-type inhibition had the most dramatic influence on low-pass filtering as evidenced by the loss of the briefer inter-event intervals. Next, we examined how the input sensitivity of TC neurons was altered by the different types of inhibition delivered at equipotent conductance levels. All three types of inhibition reduced the input sensitivity of TC neurons to all of the RGC input patterns as shown by the rightwards shift in the distribution of x_0_ values following addition of all three types of inhibition (**Figure 4E**). Therefore, across the entire population a larger change in the synaptic conductance was required to increase AP firing rates by 50%. However, the distribution of the slope coefficient (Dx) values was little changed by any type of inhibition (**Figure 4E**). By analyzing each RGC input pattern separately it was clear that the inhibitory actions were not uniform in potency across all input types. The mechanisms underlying the large variability we observed in the change in excitability using different RGC input patterns was explored in subsequent experiments. The magnitude of the conductance used to compare the impact of F1, F2 and tonic inhibition on information transfer in subsequent experiments was based on the equipotent levels of inhibition identified in Figure 4C.

**Figure 4.**
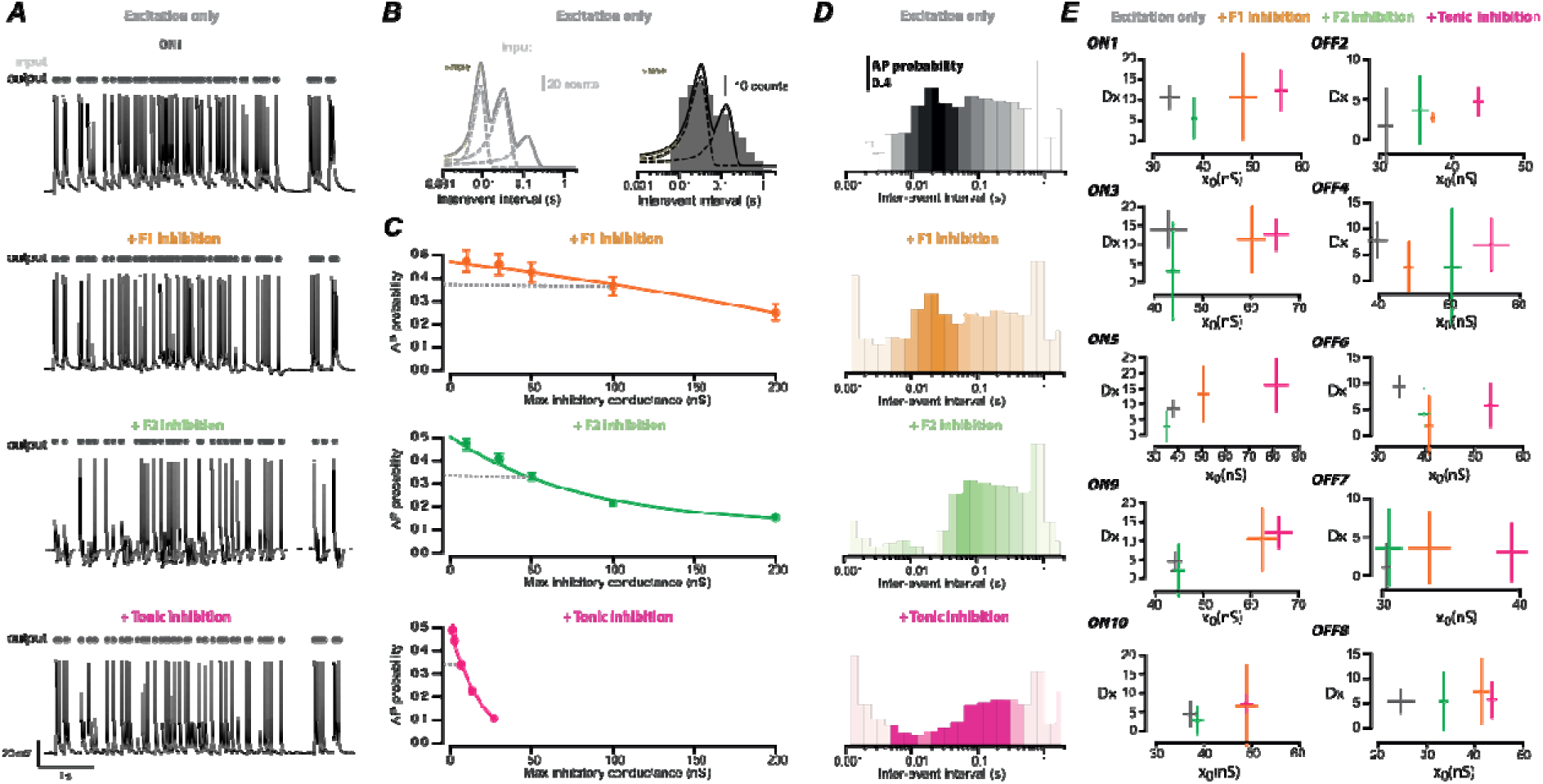
Tonic inhibition alters TC neuron input sensitivity with greater potency than phasic inhibition. *A*, Series of membrane voltage recordings obtained in response to an ON-type RGC input pattern delivered at the previously determined X_0_ value for excitation. The top trace is recorded with no inhibition present (Excitation only) whereas the next three traces include F1 inhibition, F2 inhibition and tonic inhibition that was delivered at a maximum conductance of 100, 50 & 5 nS respectively. ***B***, The left-hand histogram is an inter-event interval (IEI) distribution constructed for all 10 RGC input patterns (grey bars). The input IEI distribution was well-described by the sum of three Gaussians (smooth grey line). The right-hand histogram is an IEI distribution constructed from all AP outputs recorded from a single TC neuron (black bars). This output IEI distribution was best fit with the sum of two Gaussians (smooth black line). The shaded yellow region in each histogram denotes the high frequency region above 100 Hz. Note that for the output IEI there is a clear absence of events in this region even through these high frequency IEIs are present in the input histogram. This observation clearly illustrates the low-pass filtering characteristics of TC neurons. ***C***, Scatter plots comparing the effect of varying the maximum conductance of F1, F2 or tonic inhibition on the AP probability recorded for the cell shown in ***A***. The solid lines are the results of exponential fits to the data. The dashed line illustrates our selection of an equipotent levels of inhibition for use in subsequent experiments. ***D***, Histogram of the AP probability calculated at all IEIs following delivery of all 10 RGC input patterns to a single TC neuron. The density of the shaded bars illustrates the number of events used to calculate the AP probability at each IEI. Similar plots were constructed following the delivery F1-type, F2-type and tonic inhibition at the equipotent levels of inhibition determined from panel C. ***E***, X-Y error plots of the mean x_0_ and Dx values calculated from all TC neurons in response to each of the 10 different RGC patterns. For each RGC firing pattern, we show the input sensitivity with excitation alone and then in the presence of the equipotent levels of F1-type, F2-type and tonic inhibition that were determined in panel C. Note the consistent rightward shift in the x_0_ values that occurs with all types of inhibition is far more pronounced for tonic inhibition. However, the effect of inhibition on the slope coefficient, Dx, was far more variable and did not exhibit any obvious trends for any of the ON or OFF responses.

### The latency to first AP and rate coding are influenced in different ways by the three types of inhibition

For the next aspect of our analysis, we isolated individual clusters based upon the inter-event interval distributions of the RGC input patterns (**Figure 5A**) and examined the impact of equipotent levels of inhibition. Analysis of the latency to first AP within each cluster (**Figure 5B**) demonstrated how F1-type and F2-type inhibition enhanced AP precision compared to excitation but tonic inhibition had little impact on this aspect of information transfer (**Figure 5C**). In contrast, rate coding was shown to be more sensitive to the introduction of tonic inhibition compared to either F1-type or F2-type inhibition. The dynamic-clamp approach enabled the input sensitivity of each cluster to be estimated in the presence and absence of inhibition. As expected, all three types of inhibition tended to increase the synaptic conductance required to elicit APs in each cluster. However, tonic inhibition reduced sensitivity at all input cluster rates whereas F2-type inhibition tended not to alter the sensitivity at the lowest and highest input cluster rates (**Figure 5D**). The tonic inhibition also had the greatest impact on the input-output frequency relationship of TC neurons (Figure 5E) with AP output rates reduced across all input rates. In performing this analysis, it was noted that the clusters could be divided into two discreet populations containing low and high frequency AP rates with a clear separation at 20 s^-1^ (**Figure 5F**). For each cluster the EPSC conductance required to elicit APs within the cluster was determined and the impact of introducing inhibition was assessed on the resulting I-O relationships (**Figure 5G**). For high frequency clusters, there was a rightward shift in the I-O relationship such that a greater synaptic conductance was required to elicit APs. This change in gain was accompanied by a broader dynamic range as the slope coefficient was increased. In contrast, the action of inhibition on low frequency clusters was characterized by a leftward shift in the I-O relationship and the dynamic range became more restricted as the slope coefficient was reduced (**Figure 5H**). This effect was observed for F1-type, F2-type and tonic inhibition (**Figure 5I**) and this observation helps explain the diverse inhibitory actions we observed on Dx in earlier experiments.

**Figure 5.**
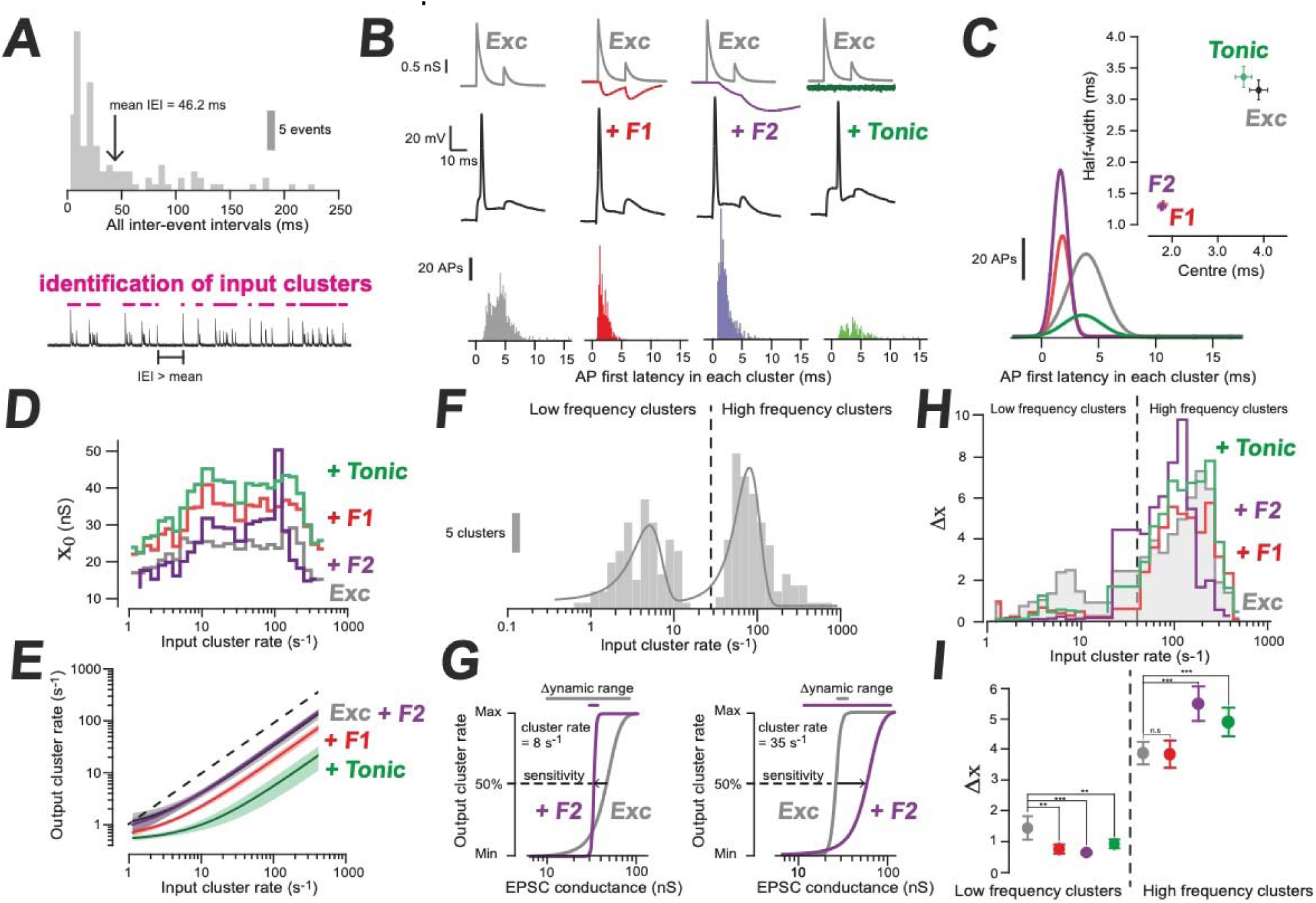
Temporal precision and rate coding are sensitive to different types of inhibition. ***A***, Histogram of the inter-event interval (IEI) distribution for an ON-type RGC input illustrating the method chosen for identifying clusters within the RGC input patterns. The mean IEI is used as the cut-off to identify each cluster shown on the conductance waveform below. In this example 13 clusters were identified. ***B***, The top traces are conductance waveforms used to excite and inhibit the TC neurons. For illustrative purposes the inhibitory waveforms are reflected downwards. The voltage traces recorded from the TC neuron are shown below each set of conductance waveforms. In each case a single AP is initiated. The distribution of AP first latency measurements obtained for the first AP initiated in each cluster are shown in the histograms. Each TC neurons received excitation from all 10 RGC input patterns and first latency distributions were constructed with excitation alone (grey), and with F1-type (red), F2-type (purple) or tonic inhibition (green). ***C***, Gaussian functions (smooth lines) were fitted to the resulting distributions obtained for each TC neuron. The scatter plot shows the mean values and SEM obtained for both the centre of each fitted gaussian and the half-width for 5 TC neurons in each condition. ***D***, Plot of the input sensitivity across all input cluster rates with excitation only (grey) and in the presence of F1-type (red), F2-type (purple) and tonic inhibition (green). ***E***, Results of exponential fits to the relationship between input frequency and the output frequency for all clusters. The shaded areas are the 95% confidence limits for the fits. ***F***, Histogram of the number of clusters at each input rate. The data has been fitted with the sum of two Gaussians. ***G***, Input-output curves for two clusters at 8 s^-1^ and 35 s^-1^. The Boltzmann functions compares the sensitivity of these clusters to changing EPSC conductance with excitation only and in the presence of F2-type inhibition. ***H***, Plot of the dynamic range across the input cluster rate for excitation only (grey) compared to addition of F1-type (red), F2-type (purple) and tonic inhibition (green). The dashed line indicates the division between low frequency and high frequency clusters as shown in ***F. I***, Comparison of the mean dynamic range for the low and high frequency clusters for excitation only (grey) compared to addition of F1-type (red), F2-type (purple) and tonic inhibition (green). ANOVA followed by Bonferroni correction was used to determine significance between response types (p<0.01: **; p<0.001: ***; n.s.: not significant).

### Information loss at the retinogeniculate synapse is minimized by tonic inhibition

Next, we examined the impact of the different forms of inhibition on the complexity of the information conveyed by the ten different RGC input patterns. The complexity of each RGC firing pattern was assessed from the Coefficient of Variability (CV_input_) and this measurement was used to assay changes in information content following AP firing in TC neurons. Ranking the input patterns based upon CV_input_ illustrated how the OFF-type inputs tended to be less complex than the ON-type RGC input patterns (**Figure 6A**). However, it was clear from a simple liner regression analysis (a slope of 0.03 and a Pearson’s r = 0.08) that the average event rate of each RGC input pattern was not the major determinant of the CV_input_. However, the more complex the RGC input pattern the more we observed a change in the CV of the TC neuron AP firing (**Figure 6B**). To examine the impact of inhibition on information loss we compared the inter-event interval distributions of the TC neuron AP output with the RGC input patterns. The difference between the input and output distributions could be assayed from the r^2^ values obtained by fitting the inter-event interval distribution of the RGC input to the AP output (**Figure 6C**). This analysis was repeated at all synaptic weights for each RGC input pattern to monitor information loss at the retinogeniculate synapse (**Figure 6D**). As expected from the previous CV analysis, F2-type inhibition resulted in the largest degree of information loss for both ON and OFF-type RGC inputs. However, this analysis did illustrate that the information loss was less following the introduction of tonic inhibition. The information loss for individual RGC inputs was then extracted and the distributions analyzed (**Figure 6E**). From these distributions we can see that the information loss was greatest for the ON-type RGC input patterns, but tonic inhibition resulted in the smallest amount of information loss. In the final analysis, based upon information loss we determined the significant difference between RGC inputs and plotted the resulting p values as heat maps (**Figure 6F**). In the presence of F1-type inhibition only two of the RGC inputs patterns resulted in AP outputs that were different from each other with ON9 showing significant differences at the p>0.05 level between both OFF2 and OFF7. F2-type inhibition was shown to maintain differences between four of the RGC input patterns while tonic inhibition was shown to maintain differences between five of the RGC input patterns. Overall, tonic inhibition provides potent gain control while maintaining information content and, therefore, preserving differences between sensory inputs to a greater extent than the other forms of gain control explored in this study.

**Figure 6.**
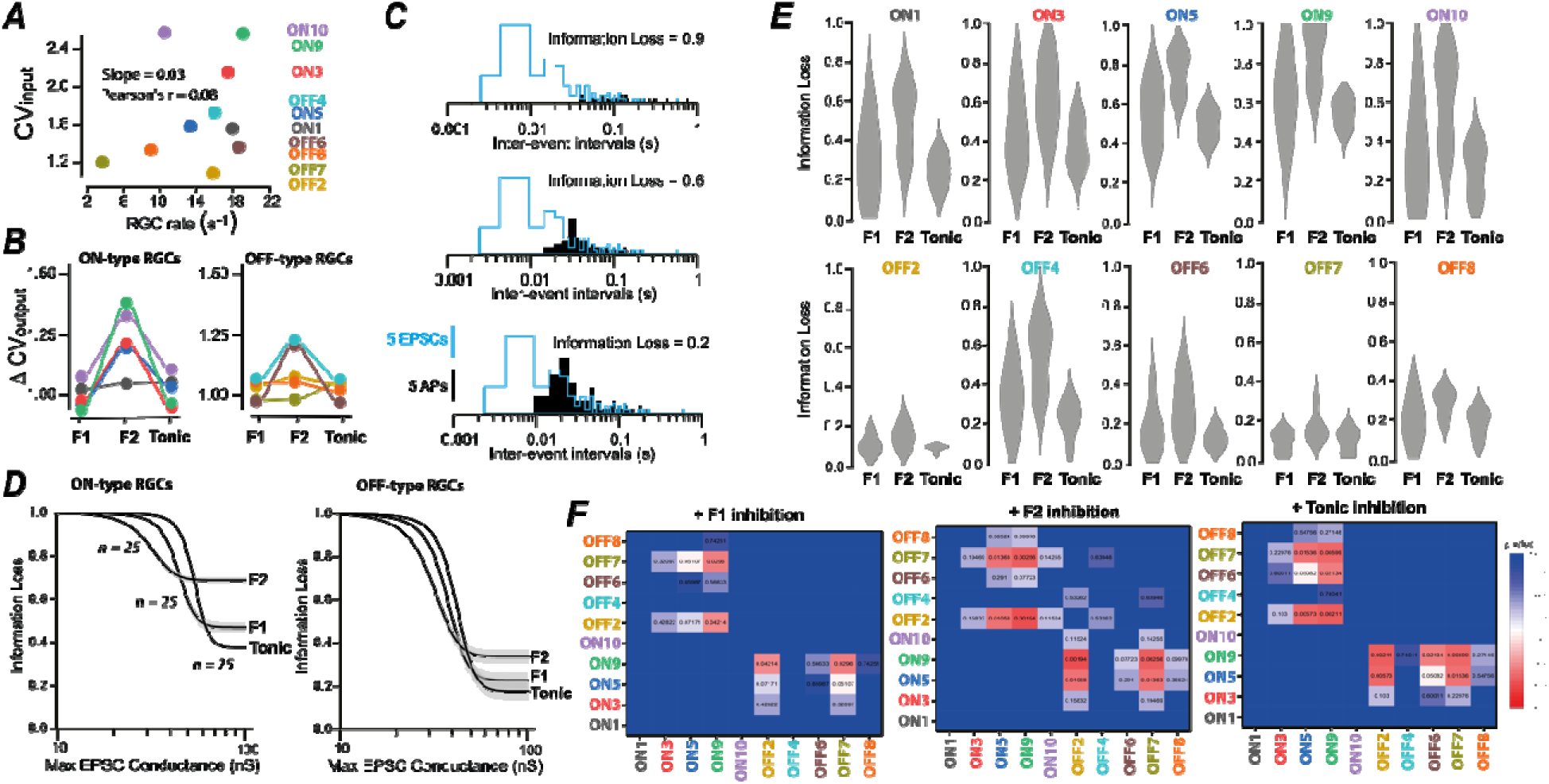
Information loss is limited by tonic inhibition. *A*, Scatter plot illustrating the relationship between the RGC event rate and the coefficient of variability (CV_input_) for each of the 10 input patterns used in this study. Linear regression analysis (not shown) indicated no relationship between these parameters with a slope of just 0.03 and a very low Pearson’s r of 0.08. ***B***, Line series plot illustrating the change in variability produced by TC neuron AP firing (CV_output_) in response to the 10 different RGC patterns following the delivery of F1, F2 and tonic inhibition at equipotent inhibition (as determined from Figure 4C). Note how F2-type inhibition was most often associated with a greater change in variability compared to F1-type and tonic inhibition. ***C,*** Histograms of inter-event intervals obtained for a single RGC input pattern (blue) superimposed on the TC neuron AP output (black) at increasing synaptic drive. The difference between the input and the output distributions was used to quantify information loss in each case. ***D,*** Plots of information loss at increasing EPSC conductance. Information loss was reduced once AP threshold was reached by the synaptic conductance and the lowest degree of information loss was consistently observed when the EPSC conductance level was at a maximum value of 100 nS. Note how the OFF-type inputs were characterised by lower levels of information loss. Consistent with previous data on CV analysis, F2-type inhibition increased the information loss whereas tonic inhibition was associated with the lowest information loss for both ON- and Off-type RGC patterns. ***E,*** A series of violin plots showing information loss for each RGC input pattern. Note how the variability was greatest in the presence of F2-type inhibition but lower for the OFF-type RGC inputs (excepting OFF4). ***F,*** Heat maps showing the significant differences that were calculated between the RGC input patterns. Note how F1-type inhibition reduces the difference between RGC-types but tonic inhibition results in the greatest number of differences between RGC input patterns.

## Discussion

This study identified clear differences in the type of GABA_A_ receptor mediated inhibition present on different TC neuron types. The X-, Y- and W-type TC neurons were first identified in the cat LGd according to the type of RGC input these morphologically distinct neurons received (Nassi & Callaway, 2009). Electron microscopy studies in the cat LGd further demonstrated that dendritic F2 terminals were presynaptic to X-type TC neurons, but F2-synapses were rarely observed onto Y-type TC neurons (Wilson *et al*., 1984; Sherman, 2004). A similar morphological classification of LGd neurons has been reported in mice (Krahe *et al*., 2011) and we now present evidence that different types of inhibitory drive are associated with these TC neuron types (**see Figure 1**). In a previous study, activation of mIPSCs by mGluRs was used to show that F2-positive relay neurons were associated with significantly slower mIPSCs, compared to F2-negative LGd relay neurons (Yang *et al*., 2017). In the current study, both averaged IPSC waveform analysis and cluster analysis revealed that reduction in interneuron density resulted in the loss of the slow-rising sIPSCs with little impact on decay times. We propose that the different clusters present in the sIPSC distributions reflect GABA_A_ receptor diversity at F1- and F2-type synapses (Ye *et al*., 2017). Combined with the morphological classification of TC neurons, our observations are consistent with the view that LGd interneuron dendrites are less likely to form F2 profiles with Y-type TC neurons (Famiglietti, 1970; Rafols & Valverde, 1973; Wilson *et al*., 1984; Hamos *et al*., 1985; Sherman, 2004) and confirm our previous reports that slow IPSCs were rarely observed onto Y-type TC neurons (Bright *et al*., 2011). Tonic inhibition has been described in many studies of TC neuron excitability (Belelli *et al*., 2005; Cope *et al*., 2005; Chandra *et al*., 2006; Bright *et al*., 2007; Bright *et al*., 2011; Herd *et al*., 2013) and we have shown that tonic inhibition is a feature of both X- and Y-type TC neurons (Bright *et al*., 2007; Bright *et al*., 2011). The dynamic-clamp experiments employed in this study (**see below**) demonstrate how the different types of inhibition present in the thalamus have distinct influences on various aspects of information transfer by thalamocortical neurons.

Our dynamic-clamp protocols were designed to mimic the fuzzy logic associated with RGC input onto TC neurons in the LGd (Morgan *et al*., 2016; Rompani *et al*., 2017) while delivering equipotent levels (see **Figure 4C**) of F1-type, F2-type, and tonic inhibition. This level of control would not have been feasible with pharmacological manipulations or any conceivable optogenetic approach. Using this approach, we were able to demonstrate how the input sensitivity of TC neurons to 10 different RGC input patterns was remarkably similar in the absence of inhibition (see **Figure 3I**). The input-output relationships revealed very similar x_0_ and Dx values in response to the 10 different RGC input patterns. At the same time each TC neuron was able to convey the information conveyed by these different RGC inputs as distinct AP output firing patterns (see **Figure 3H**). This result demonstrates how a TC neuron can discriminate between the information conveyed by different RGC inputs independent of the weighing of each retinogeniculate synaptic input (Litvina & Chen, 2017). Importantly, we show how the synchronous GABA release associated with F1-type inhibition degrades the ability of a TC neuron to distinguish between different RGC inputs to a much greater extent than the asynchronous GABA release associated with either F2-type or tonic inhibition (see **Figure 6F**). These observations offer a compelling insight into why non-vesicular GABA release has evolved to become a dominant influence on TC excitability within a first order sensory thalamic region like the LGd where information contained within RGC firing patterns needs to be faithfully conveyed to the neocortex. It is intriguing to speculate that by varying the contribution of these three different types of inhibition the thalamus can alter input selectivity without the need to alter synaptic weighting of the retinogeniculate input. Importantly, this computational flexibility will not be associated with the metabolic demands of altering synaptic weighting.

One obvious shortcoming of this study is our assumption that LGd interneurons can faithfully convey the information arriving from the different RGC types into unique patterns of inhibition. Unfortunately, the efficacy of feed-forward phasic inhibition generated by the RGC-to-interneuron-to-TC circuit has yet to be adequately studied. However, calcium influx into the dendrites of LGd interneurons has been shown to elicit asynchronous GABA release onto TC neurons (Acuna-Goycolea *et al*., 2008) indicating how temporal information may be lost during feedforward inhibition of TC neurons. In our study, we have also shown that optogenetic GABA release from LGd interneurons generates delayed postsynaptic responses onto TC neurons (see **Figure 2**). Notably, our previous studies failed to demonstrate reliable feedforward phasic inhibition during simultaneous recording from LGd interneurons and TC neurons (Jager *et al*., 2016). Similarly, in the somatosensory thalamus, simultaneous recording between 260 pairs of local interneurons and TC neurons resulted in only 9 clear examples of AP-dependent functional connectivity (Simko & Markram, 2021). It is worth considering that identification of functional connectivity in these paired recording studies was based upon the coincidence of presynaptic APs with postsynaptic changes, and it is conceivable that a greater level of functional connectivity would have been reported if sub-threshold dendro-dendritic GABA release was assayed. Therefore, our future studies will focus on examining the importance of sub-threshold voltage changes in the dendrites of LGd interneurons, elicited by RGC input, for controlling the timing of TC inhibition. Another limitation of this study is our focus on a single RGC inputs ability to determine the pattern of inhibition generated by a local interneuron at F1 and F2 type synapses onto TC neurons. There is of course compelling evidence that a local LGd interneuron receives afferent input from multiple RGC inputs and that the inhibitory output onto TC neurons could, therefore, reflect summation from many RGC types (Morgan & Lichtman, 2020). This has been demonstrated functionally by recording from local interneurons during electrical stimulation of the optic tract (Seabrook *et al*., 2013).. In future studies, we plan to generate patterns of inhibition that reflect different levels of RGC summation but at present the rules of this connectivity and the transformation properties of the local interneurons are far from clear.

It has yet to be determined whether the lagged inhibition associated with F2-type dendro-dendritic synapses reflects (1) a noncanonical form of exocytosis (Kennedy & Ehlers, 2011) associated with GABA release from LGd interneuron dendrites (Acuna-Goycolea *et al*., 2008), (2) GABA spillover onto distant GABA_A_ receptors located within a glomerular structure (Rossi & Hamann, 1998), or (3) the presence of GABA_A_ receptor types with particularly slow activation kinetics (Ye *et al*., 2017). Whatever the etiology of the slow-rising IPSCs it is clear from our dynamic-clamp experiments that F2-type inhibition has a distinct influence on information transfer compared to F1-type and tonic inhibition. The experiments undertaken in this study demonstrate how both F1- and F2-type feed-forward inhibition onto TC neurons will enhance temporal precision whereas all three types of inhibition have a similar influence on rate coding within the thalamus (see **Figure 5**).

We have previously reported how high interneuron AP firing rates generate tonic inhibition onto TC neurons due to the persistent activation of high-affinity extrasynaptic δ-GABA_A_Rs by ambient GABA levels in the extracellular space (Jager *et al*., 2016). The asynchronous and lagged response of dendro-dendritic F2-type inhibition will further contribute to the appearance of tonic inhibition onto TC neurons at high input frequencies. Raised ambient GABA levels are best explained by the stoichiometry of the GAT-1 transporter (Wu *et al*., 2007) but may also be associated with GABA release from astrocytes through bestrophin channels (Lee *et al*., 2010). Irrespective of the non-vesicular source of GABA, our results are consistent with observations made in the VB that tactile discrimination is improved by enhanced tonic inhibition (Kwak *et al*., 2020) and supports a long-standing biophysical model that has explained how tonic inhibition improves neuronal precision due to alterations in the membrane time constant (Semyanov *et al*., 2004).

The hypothesis that increasing numbers of local GABAergic interneurons within thalamic circuits reflects a requirement for more complex levels of neural processing was first proposed by Cajal (Cajal, 1909; Cajal & Ramón-Moliner, 1966) and more recent immunocytochemical data has shown how the (Annecchino *et al*., 2017) proportion of these interneurons progressively increases in species associated with more complex behavior (Arcelli *et al*., 1997). Contrary to earlier studies (Barbaresi *et al*., 1986; Smith *et al*., 1987; Bentivoglio *et al*., 1991) it is now apparent that local interneurons are present in the dorsal thalamic nuclei of both primates and rodents (Jager *et al*., 2021) but interneuron numbers are particularly enriched in the LGd of many mammalian species including rodents (Khan *et al*., 1994; Arcelli *et al*., 1997). The presence of local interneurons specifically within the rodent LGd could be a consequence of greater information content travelling through the visual thalamus or may reflect a requirement for additional types of sensory processing. It could be argued that the former may not require a change in the nature of the inhibitory control but, the latter may introduce a pressure to diversify the characteristics of inhibition. However, our study demonstrates that thalamic interneurons have the potential to influence sensory processing in unique and unexpected ways depending on the nature of GABA_A_ receptor activation. The computational flexibility afforded by altering input selectivity by simply varying the contribution of synchronous and asynchronous GABA release is a model that is likely to be observed beyond the visual thalamus.

## Reference

Acuna-Goycolea, C., Brenowitz, S.D. & Regehr, W.G. (2008) Active dendritic conductances dynamically regulate GABA release from thalamic Interneurons. Neuron, 57, 420–431.

Annecchino, L.A., Morris, A.R., Copeland, C.S., Agabi, O.E., Chadderton, P. & Schultz, S.R. (2017) Robotic Automation of In Vivo Two-Photon Targeted Whole-Cell Patch-Clamp Electrophysiology. Neuron, 95, 1048–1055 e1043.

Arcelli, P., Frassoni, C., Regondi, M.C., De Biasi, S. & Spreafico, R. (1997) GABAergic neurons in mammalian thalamus: a marker of thalamic complexity? Brain research bulletin, 42, 27–37.

Bakken, T.E., van Velthoven, C.T., Menon, V., Hodge, R.D., Yao, Z., Nguyen, T.N., Graybuck, L.T., Horwitz, G.D., Bertagnolli, D., Goldy, J., Yanny, A.M., Garren, E., Parry, S., Casper, T., Shehata, S.I., Barkan, E.R., Szafer, A., Levi, B.P., Dee, N., Smith, K.A., Sunkin, S.M., Bernard, A., Phillips, J., Hawrylycz, M.J., Koch, C., Murphy, G.J., Lein, E., Zeng, H. & Tasic, B. (2021) Single-cell and single-nucleus RNA-seq uncovers shared and distinct axes of variation in dorsal LGN neurons in mice, non-human primates, and humans. Elife, 10.

Barbaresi, P., Spreafico, R., Frassoni, C. & Rustioni, A. (1986) GABAergic neurons are present in the dorsal column nuclei but not in the ventroposterior complex of rats. Brain Res, 382, 305–326.

Belelli, D., Peden, D.R., Rosahl, T.W., Wafford, K.A. & Lambert, J.J. (2005) Extrasynaptic GABAA receptors of thalamocortical neurons: a molecular target for hypnotics. The Journal of neuroscience: the official journal of the Society for Neuroscience, 25, 11513–11520.

Bentivoglio, M., Spreafico, R., Minciacchi, D. & Macchi, G. (1991) GABAergic interneurons and neuropil of the intralaminar thalamus: an immunohistochemical study in the rat and the cat, with notes in the monkey. Exp Brain Res, 87, 85–95.

Bishop, P.O., Burke, W. & Davis, R. (1958) Synapse discharge by single fibre in mammalian visual system. Nature, 182, 728–730.

Blitz, D.M. & Regehr, W.G. (2003) Retinogeniculate synaptic properties controlling spike number and timing in relay neurons. Journal of neurophysiology, 90, 2438–2450.

Blitz, D.M. & Regehr, W.G. (2005) Timing and Specificity of Feed-Forward Inhibition within the LGN. Neuron, 45, 917–928.

Brickley, S.G. & Mody, I. (2012) Extrasynaptic GABA(A) receptors: their function in the CNS and implications for disease. Neuron, 73, 23–34.

Bright, D.P., Aller, M.I. & Brickley, S.G. (2007) Synaptic release generates a tonic GABA(A) receptor-mediated conductance that modulates burst precision in thalamic relay neurons. The Journal of neuroscience: the official journal of the Society for Neuroscience, 27, 2560–2569.

Bright, D.P., Renzi, M., Bartram, J., McGee, T.P., MacKenzie, G., Hosie, A.M., Farrant, M. & Brickley, S.G. (2011) Profound desensitization by ambient GABA limits activation of delta-containing GABAA receptors during spillover. The Journal of neuroscience: the official journal of the Society for Neuroscience, 31, 753–763.

Brock, O., Gelegen, C., Sully, P., Salgarella, I., Jager, P., Menage, L., Mehta, I., Jeczmien-Lazur, J., Djama, D., Strother, L., Coculla, A., Vernon, A.C., Brickley, S., Holland, P., Cooke, S.F. & Delogu, A. (2022) A Role for Thalamic Projection GABAergic Neurons in Circadian Responses to Light. The Journal of neuroscience: the official journal of the Society for Neuroscience, 42, 9158–9179.

Cajal, S.R. (1909) Histologie du système nerveux de l’homme & des vertébrés. Paris: Maloine.

Cajal, S.R. & Ramón-Moliner, E. (1966) Studies on the diencephalon. C. C. Thomas, Springfield, Ill.,.

Casale, A.E. & McCormick, D.A. (2011) Active action potential propagation but not initiation in thalamic interneuron dendrites. The Journal of neuroscience: the official journal of the Society for Neuroscience, 31, 18289–18302.

Chandra, D., Jia, F., Liang, J., Peng, Z., Suryanarayanan, A., Werner, D.F., Spigelman, I., Houser, C.R., Olsen, R.W., Harrison, N.L. & Homanics, G.E. (2006) GABAA receptor alpha 4 subunits mediate extrasynaptic inhibition in thalamus and dentate gyrus and the action of gaboxadol. Proc Natl Acad Sci U S A, 103, 15230–15235.

Chen, C., Blitz, D.M. & Regehr, W.G. (2002) Contributions of receptor desensitization and saturation to plasticity at the retinogeniculate synapse. Neuron, 33, 779–788.

Cleland, B.G., Dubin, M.W. & Levick, W.R. (1971) Simultaneous recording of input and output of lateral geniculate neurones. Nat New Biol, 231, 191–192.

Cope, D.W., Hughes, S.W. & Crunelli, V. (2005) GABAA receptor-mediated tonic inhibition in thalamic neurons. The Journal of neuroscience: the official journal of the Society for Neuroscience, 25, 11553–11563.

Crandall, S.R. & Cox, C.L. (2012) Local dendrodendritic inhibition regulates fast synaptic transmission in visual thalamus. The Journal of neuroscience: the official journal of the Society for Neuroscience, 32, 2513–2522.

Crone, S.A., Quinlan, K.A., Zagoraiou, L., Droho, S., Restrepo, C.E., Lundfald, L., Endo, T., Setlak, J., Jessell, T.M., Kiehn, O. & Sharma, K. (2008) Genetic ablation of V2a ipsilateral interneurons disrupts left-right locomotor coordination in mammalian spinal cord. Neuron, 60, 70–83.

Delogu, A., Sellers, K., Zagoraiou, L., Bocianowska-Zbrog, A., Mandal, S., Guimera, J., Rubenstein, J.L., Sugden, D., Jessell, T. & Lumsden, A. (2012) Subcortical visual shell nuclei targeted by ipRGCs develop from a Sox14+-GABAergic progenitor and require Sox14 to regulate daily activity rhythms. Neuron, 75, 648–662.

Famiglietti, E.V., Jr. (1970) Dendro-dendritic synapses in the lateral geniculate nucleus of the cat. Brain Res, 20, 181–191.

Ferraguti, F., Cobden, P., Pollard, M., Cope, D., Shigemoto, R., Watanabe, M. & Somogyi, P. (2004) Immunolocalization of metabotropic glutamate receptor 1alpha (mGluR1alpha) in distinct classes of interneuron in the CA1 region of the rat hippocampus. Hippocampus, 14, 193–215.

Fiala, J.C. (2005) Reconstruct: a free editor for serial section microscopy. Journal of microscopy, 218, 52–61.

Friedlander, M.J., Lin, C.S., Stanford, L.R. & Sherman, S.M. (1981) Morphology of functionally identified neurons in lateral geniculate nucleus of the cat. Journal of neurophysiology, 46, 80–129.

Grubb, M.S. & Thompson, I.D. (2003) Quantitative characterization of visual response properties in the mouse dorsal lateral geniculate nucleus. Journal of neurophysiology, 90, 3594–3607.

Hammer, S., Monavarfeshani, A., Lemon, T., Su, J. & Fox, M.A. (2015) Multiple Retinal Axons Converge onto Relay Cells in the Adult Mouse Thalamus. Cell Rep, 12, 1575–1583.

Hamos, J.E., Van Horn, S.C., Raczkowski, D., Uhlrich, D.J. & Sherman, S.M. (1985) Synaptic connectivity of a local circuit neurone in lateral geniculate nucleus of the cat. Nature, 317, 618–621.

Harris, J.J., Jolivet, R., Engl, E. & Attwell, D. (2015) Energy-Efficient Information Transfer by Visual Pathway Synapses. Curr Biol, 25, 3151–3160.

Herd, M.B., Brown, A.R., Lambert, J.J. & Belelli, D. (2013) Extrasynaptic GABA(A) receptors couple presynaptic activity to postsynaptic inhibition in the somatosensory thalamus. The Journal of neuroscience: the official journal of the Society for Neuroscience, 33, 14850–14868.

Houston, C.M., Bright, D.P., Sivilotti, L.G., Beato, M. & Smart, T.G. (2009) Intracellular chloride ions regulate the time course of GABA-mediated inhibitory synaptic transmission. The Journal of neuroscience: the official journal of the Society for Neuroscience, 29, 10416–10423.

Jager, P., Moore, G., Calpin, P., Durmishi, X., Salgarella, I., Menage, L., Kita, Y., Wang, Y., Kim, D.W., Blackshaw, S., Schultz, S.R., Brickley, S., Shimogori, T. & Delogu, A. (2021) Dual midbrain and forebrain origins of thalamic inhibitory interneurons. Elife, 10.

Jager, P., Ye, Z., Yu, X., Zagoraiou, L., Prekop, H.T., Partanen, J., Jessell, T.M., Wisden, W., Brickley, S.G. & Delogu, A. (2016) Tectal-derived interneurons contribute to phasic and tonic inhibition in the visual thalamus. Nature communications, 7, 13579.

Kennedy, M.J. & Ehlers, M.D. (2011) Mechanisms and function of dendritic exocytosis. Neuron, 69, 856–875.

Khan, A.A., Wadhwa, S. & Bijlani, V. (1994) Development of human lateral geniculate nucleus: an electron microscopic study. Int J Dev Neurosci, 12, 661–672.

Krahe, T.E., El-Danaf, R.N., Dilger, E.K., Henderson, S.C. & Guido, W. (2011) Morphologically distinct classes of relay cells exhibit regional preferences in the dorsal lateral geniculate nucleus of the mouse. The Journal of neuroscience: the official journal of the Society for Neuroscience, 31, 17437–17448.

Kwak, H., Koh, W., Kim, S., Song, K., Shin, J.I., Lee, J.M., Lee, E.H., Bae, J.Y., Ha, G.E., Oh, J.E., Park, Y.M., Kim, S., Feng, J., Lee, S.E., Choi, J.W., Kim, K.H., Kim, Y.S., Woo, J., Lee, D., Son, T., Kwon, S.W., Park, K.D., Yoon, B.E., Lee, J., Li, Y., Lee, H., Bae, Y.C., Lee, C.J. & Cheong, E. (2020) Astrocytes Control Sensory Acuity via Tonic Inhibition in the Thalamus. Neuron, 108, 691–706 e610.

Lee, S., Yoon, B.E., Berglund, K., Oh, S.J., Park, H., Shin, H.S., Augustine, G.J. & Lee, C.J. (2010) Channel-mediated tonic GABA release from glia. Science, 330, 790–796.

Litvina, E.Y. & Chen, C. (2017) Functional Convergence at the Retinogeniculate Synapse. Neuron, 96, 330–338 e335.

Llinas, R. & Jahnsen, H. (1982) Electrophysiology of mammalian thalamic neurones in vitro. Nature, 297, 406–408.

Mazzoni, A., Broccard, F.D., Garcia-Perez, E., Bonifazi, P., Ruaro, M.E. & Torre, V. (2007) On the dynamics of the spontaneous activity in neuronal networks. PLoS One, 2, e439.

Meytlis, M., Nichols, Z. & Nirenberg, S. (2012) Determining the role of correlated firing in large populations of neurons using white noise and natural scene stimuli. Vision Res, 70, 44–53.

Montero, V.M. & Scott, G.L. (1981) Synaptic terminals in the dorsal lateral geniculate nucleus from neurons of the thalamic reticular nucleus: a light and electron microscope autoradiographic study. Neuroscience, 6, 2561–2577.

Morgan, J.L., Berger, D.R., Wetzel, A.W. & Lichtman, J.W. (2016) The Fuzzy Logic of Network Connectivity in Mouse Visual Thalamus. Cell, 165, 192–206.

Morgan, J.L. & Lichtman, J.W. (2020) An Individual Interneuron Participates in Many Kinds of Inhibition and Innervates Much of the Mouse Visual Thalamus. Neuron, 106, 468–481 e462.

Nassi, J.J. & Callaway, E.M. (2009) Parallel processing strategies of the primate visual system. Nat Rev Neurosci, 10, 360–372.

Ohara, P.T., Lieberman, A.R., Hunt, S.P. & Wu, J.Y. (1983) Neural elements containing glutamic acid decarboxylase (GAD) in the dorsal lateral geniculate nucleus of the rat; immunohistochemical studies by light and electron microscopy. Neuroscience, 8, 189–211.

Rafols, J.A. & Valverde, F. (1973) The structure of the dorsal lateral geniculate nucleus in the mouse. A Golgi and electron microscopic study. The Journal of comparative neurology, 150, 303–332.

Ragan, T., Kadiri, L.R., Venkataraju, K.U., Bahlmann, K., Sutin, J., Taranda, J., Arganda-Carreras, I., Kim, Y., Seung, H.S. & Osten, P. (2012) Serial two-photon tomography for automated ex vivo mouse brain imaging. Nat Methods, 9, 255–258.

Rompani, S.B., Mullner, F.E., Wanner, A., Zhang, C., Roth, C.N., Yonehara, K. & Roska, B. (2017) Different Modes of Visual Integration in the Lateral Geniculate Nucleus Revealed by Single-Cell-Initiated Transsynaptic Tracing. Neuron, 93, 1519.

Rossi, D.J. & Hamann, M. (1998) Spillover-mediated transmission at inhibitory synapses promoted by high affinity alpha6 subunit GABA(A) receptors and glomerular geometry. Neuron, 20, 783–795.

Seabrook, T.A., Krahe, T.E., Govindaiah, G. & Guido, W. (2013) Interneurons in the mouse visual thalamus maintain a high degree of retinal convergence throughout postnatal development. Neural development, 8, 24.

Semyanov, A., Walker, M.C., Kullmann, D.M. & Silver, R.A. (2004) Tonically active GABA A receptors: modulating gain and maintaining the tone. Trends in neurosciences, 27, 262–269.

Sherman, S.M. (2001) Thalamic relay functions. Prog Brain Res, 134, 51–69.

Sherman, S.M. (2004) Interneurons and triadic circuitry of the thalamus. Trends in neurosciences, 27, 670–675.

Simko, J. & Markram, H. (2021) Morphology, physiology and synaptic connectivity of local interneurons in the mouse somatosensory thalamus. J Physiol, 599, 5085–5101.

Smith, Y., Seguela, P. & Parent, A. (1987) Distribution of GABA-immunoreactive neurons in the thalamus of the squirrel monkey (Saimiri sciureus). Neuroscience, 22, 579–591.

Sumitomo, I., Nakamura, M. & Iwama, K. (1976) Location and function of the so-called interneurons of rat lateral geniculate body. Experimental neurology, 51, 110–123.

Usrey, W.M., Reppas, J.B. & Reid, R.C. (1999) Specificity and strength of retinogeniculate connections. Journal of neurophysiology, 82, 3527–3540.

Wilson, J.R., Friedlander, M.J. & Sherman, S.M. (1984) Fine structural morphology of identified X- and Y-cells in the cat’s lateral geniculate nucleus. Proc R Soc Lond B Biol Sci, 221, 411–436.

Wu, Y., Wang, W., Diez-Sampedro, A. & Richerson, G.B. (2007) Nonvesicular inhibitory neurotransmission via reversal of the GABA transporter GAT-1. Neuron, 56, 851–865.

Xu, W. & Baker, S.N. (2018) In vitro characterization of intrinsic properties and local synaptic inputs to pyramidal neurons in macaque primary motor cortex. Eur J Neurosci, 48, 2071–2083.

Yang, S., Govindaiah, G., Lee, S.H., Yang, S. & Cox, C.L. (2017) Distinct kinetics of inhibitory currents in thalamocortical neurons that arise from dendritic or axonal origin. PLoS One, 12, e0189690.

Ye, Z., Yu, X., Houston, C.M., Aboukhalil, Z., Franks, N.P., Wisden, W. & Brickley, S.G. (2017) Fast and Slow Inhibition in the Visual Thalamus Is Influenced by Allocating GABAA Receptors with Different γ Subunits. Frontiers in Cellular Neuroscience, 11.

